# MALDI Tandem Mass Spectrometry for Colony-Based Dereplication of Natural Products

**DOI:** 10.64898/2026.06.21.733640

**Authors:** Robert A. Shepherd, Limar Y. Gad, Michael Strobel, Gordon T. Luu, Jin Feng, Carlo De Silva, Shaun M.K. McKinnie, Mingxun Wang, Laura M. Sanchez

## Abstract

Microbial libraries remain an important resource for natural product discovery; however, constructing taxonomically and chemically diverse collections remains a challenge. Advances in dereplication strategies, including molecular networking, have reduced the rediscovery of known bioactive molecules and facilitated the identification of novel chemical scaffolds, but these approaches are typically applied after library construction or to existing repositories. Furthermore, many dereplication workflows require scaled fermentation and extraction, increasing the time needed to assess a microbe’s metabolite profile. Here, we integrate matrix-assisted laser desorption/ionization tandem mass spectrometry (MALDI-MS/MS) into the bioinformatics platform IDBac, enabling streamlined characterization of microbial taxonomic identity, metabolite production potential, and preliminary metabolite annotation through GNPS2 molecular networking. This miniaturized high-content workflow facilitates strain prioritization by providing metabolite annotations directly from single microbial colonies prior to scale-up and extraction. Application of this approach to marine actinomycetes enabled the annotation of lavanducyanin and multiple napyradiomycin analogs. Subsequent investigation led to the discovery of napyradiomycin B8 from marine *Streptomyces* sp. CNZ-289, which was confirmed by 1D and 2D NMR spectroscopy and MALDI-MS/MS. Expanding this workflow to an untargeted analysis of 25 commensal marine vertebrate-derived bacterial isolates resulted in the annotation of several known bioactive natural products, including surugamides, antimycins, desferrioxamine siderophores, and the isolation and elucidation of harmane derivatives using NMR.

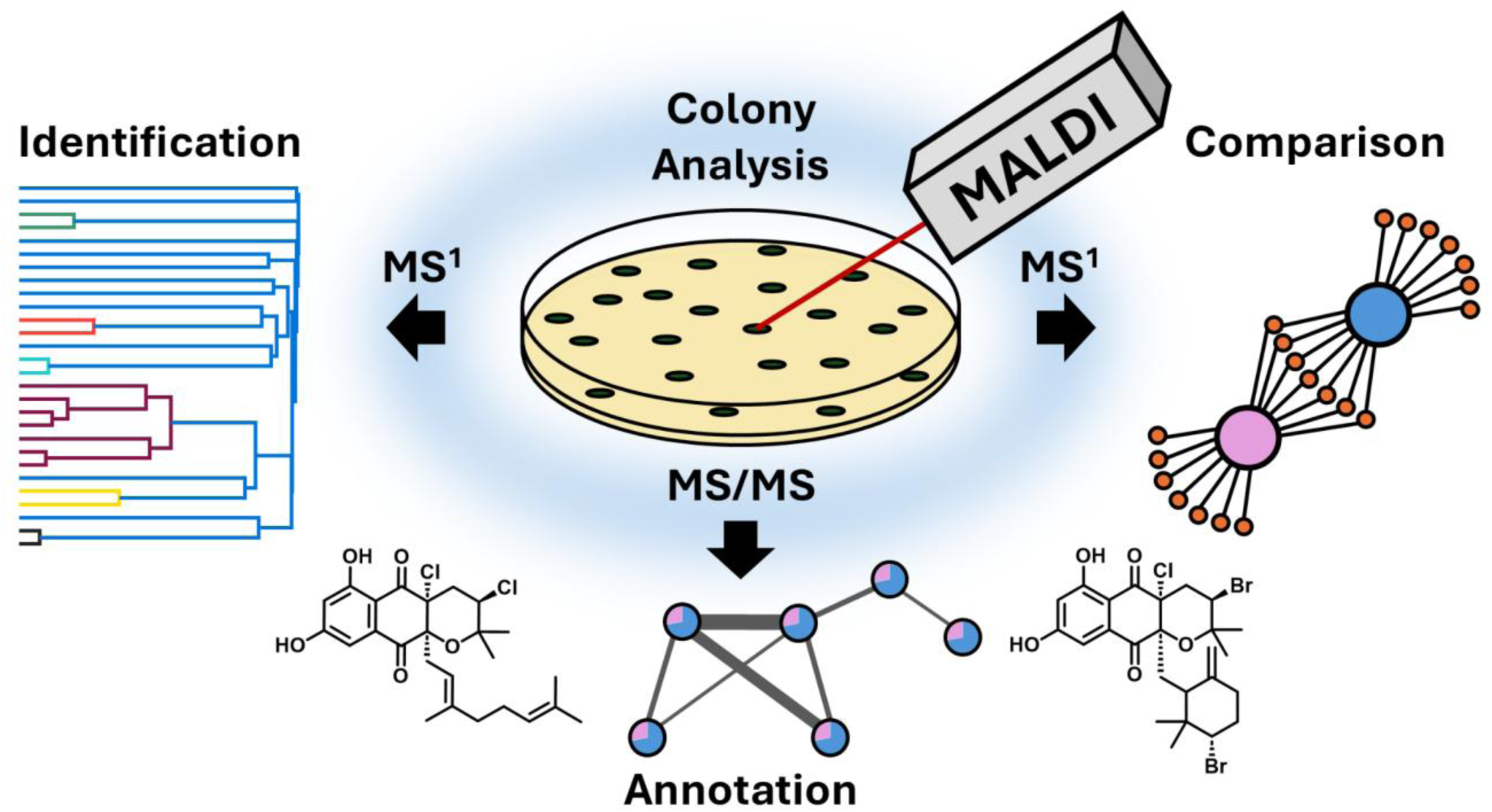

## Introduction

Microbes represent a unique reservoir for chemically diverse small molecules, otherwise known as natural products or specialized metabolites.^1^ Microbial natural products have been a cornerstone in drug development since the identification of penicillin ushered in the “Golden Age” of antibiotic discovery.^2,3^ This led to widespread construction of vast microbial libraries to mine for bioactive natural products. However, while useful these libraries often contained various replicates across species which has contributed to high rates of rediscovery. To improve decision-making during microbial selection from environmental plates, IDBac was developed as a high-throughput bioinformatics platform. By integrating matrix-assisted laser desorption/ionization-time-of-flight (MALDI-TOF) protein profiles with specialized metabolite spectra, IDBac rapidly differentiates bacterial species and links microbial identity to small-molecule production potential through direct analysis of colony biomass.^4^

At the protein level, IDBac leverages high-copy number ribosomal proteins in MALDI-TOF spectra gathered from 2-20 kDa as “fingerprints” associated with microbial identity. Construction of dendrograms via hierarchical clustering of protein spectral similarity organizes isolates into putative taxonomic groupings.^5–9^ IDBac differentiates itself from other MALDI-based microbial identification platforms^10–12^ by implementing MS spectra of specialized metabolite profiles and allowing their visualization through metabolite association networks (MANs). IDBac can discriminate closely related bacterial isolates (indistinguishable by 16S rRNA sequence) based on inter- and intraspecies differences in specialized metabolite production via the small molecule MS spectra (100-2000 Da).^4^ As a proof of principle, IDBac was reported to reduce an existing library of 833 microbial isolates down to 233 isolates (72.0% reduction) on the basis of protein similarity and metabolic overlap between specialized metabolite MS spectra of bacterial colonies.^5^ The IDBac pipeline was recently introduced as an open-access web tool and knowledgebase (KB) within the GNPS2 environment, expanding access to the broader natural products community.^13^

One limitation of the original IDBac platform is the limited structural information obtained from MALDI-MS^1^, which restricts the ability to effectively dereplicate isolates based on the detected natural product identity. Although liquid chromatography-tandem mass spectrometry (LC-MS/MS) can, and should, be used to orthogonally to validate metabolites identified during IDBac analyses, this typically occurs post-prioritization and often requires scale-up, extraction, and extended data acquisition. Herein, we address this limitation by integrating MALDI data dependent acquisition (DDA) into the existing IDBac pipeline, enabling natural product annotation directly from single microbial colonies prior to scale-up and extraction. MALDI-MS/MS data can now be deposited alongside MALDI protein and small molecule MS data, facilitating downstream molecular networking within the IDBac x GNPS2 environment.

We tested this new pipeline on a small set of known Actinomycete strains and identified lavanducyanin, and napyradiomycins A1 and B3 via MALDI-MS/MS molecular networking analysis. Further investigations led to the discovery of a new napyradiomycin B analogue whose structure was verified using MALDI-MS/MS alongside 1D and 2D NMR experiments. Further, 25 microbial strains that had been previously isolated from marine vertebrate intestines^14,15^ were queried and prioritized for scale up based on the following three criteria: 1) strains appeared taxonomically unique within this collection, 2) demonstrated production potential as measured by unique features within the MANs, and 3) appeared to have unique clusters within a molecular network. Applying this logic resulted in the dereplication of surfactins B and C, surugamide A, and antimycin A1 directly from the colonies of various marine-derived bacteria without the need for chromatography. Utilization of this IDBac x GNPS2 workflow enabled sample preparation, protein and small molecule MALDI-MS^1^ acquisition, small molecule MALDI-MS/MS acquisition, and data analysis to be completed in approximately 8 hours. As a result, researchers can obtain putative microbial identity, associated metabolite profiles, and preliminary small-molecule annotations within a single workday, maximizing the information available prior to microbial scale-up and extraction. It should be noted that while we identified and isolated known compounds using MS/MS and NMR analyses, some did not have spectra within known databases, highlighting a persistent need to grow these databases with user submissions.

## Results and Discussion

### Workflow Optimization

In order to collect data, it was first necessary to develop a strategy for automatically collecting data-dependent acquisition (DDA) on the timsTOF fleX mass spectrometer. While older MALDI-TOF based instruments had this capability, it has been removed from newer software platforms required for data collection. To address this, we developed a Python script that would allow for the selection of precursor *m/z* values directly from the MALDI-MS^1^ files used for the construction of MANs in the IDBac web platform. The script provides a graphical user interface (GUI) that allows the user to preprocess MALDI-MS^1^ data, filter and exclude MALDI matrix signals, and generate *.run files containing a list of precursor *m/z* values to be fragmented (See **Supporting Information** for more details). The *.run files can be loaded directly into timsControl for MS/MS acquisition. It is important to note that this approach utilizes “offline” MS/MS acquisition, and is not an “online” DDA technique as is the standard for LC-MS/MS. The full analysis workflow is performed as follows: (1) direct colony transfer and MALDI matrix application; (2) MALDI-MS^1^ acquisition within the small molecule (100-2000 Da) and protein (2-20 kDa) mass ranges, followed by data visualization with IDBac via MANs and protein dendrograms; and (3) generation of MS/MS precursor lists from the MALDI-MS^1^ small molecule data to perform MALDI-MS/MS and downstream molecular networking in GNPS2 (**Figure 1**).

**Figure 1.**
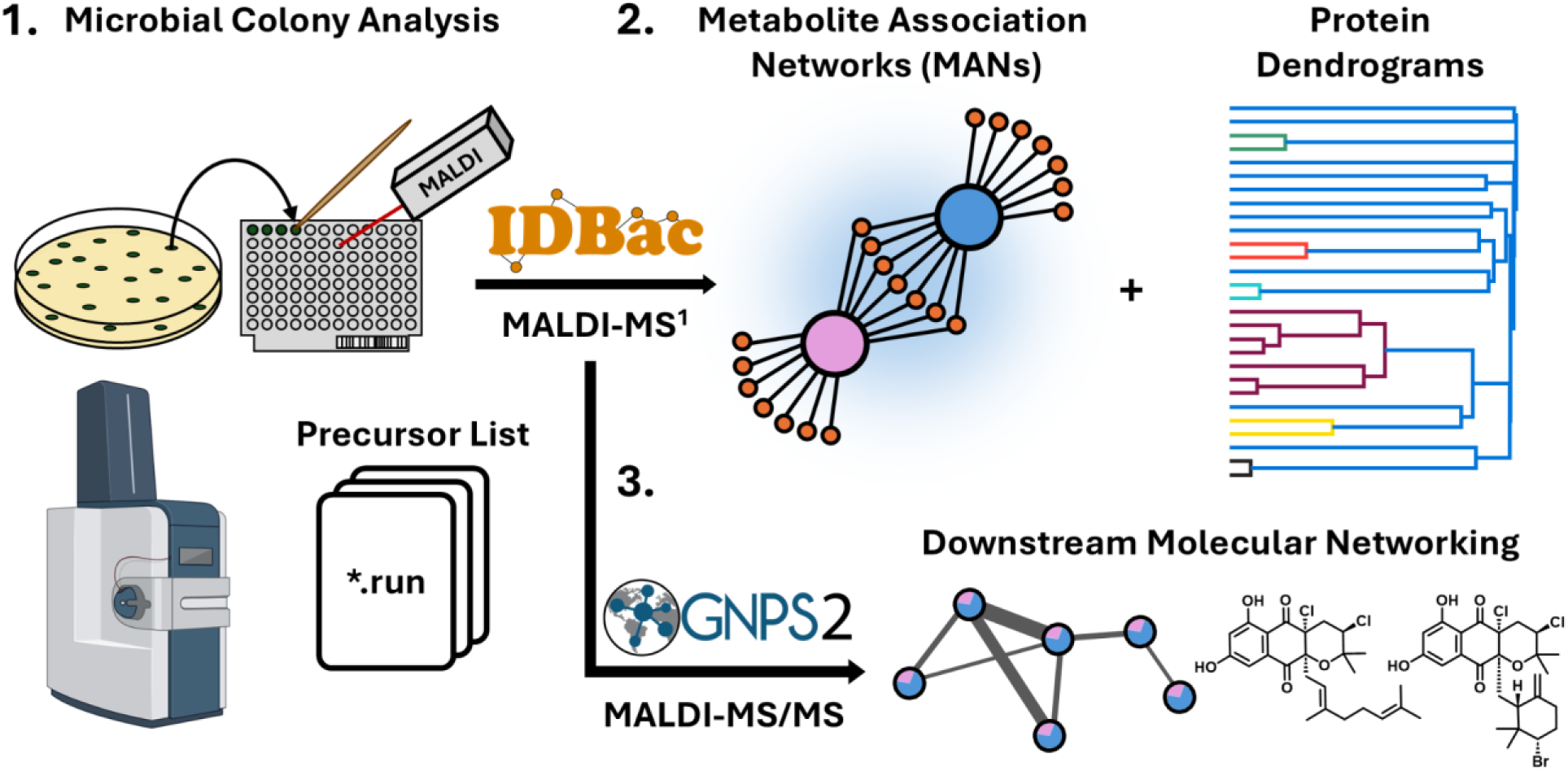
Stepwise schematic of the MALDI-DDA workflow used for the collection of MALDI-MS/MS data in this study that facilitates analysis in the GNPS2 x IDBac ecosystem. Logos have been adapted and reprinted with permission from the creators.

Next we sought to optimize the choice of organic matrix for MALDI-DDA analysis, as this can have a direct effect on the ionization of certain molecular species.^16–18^ We tested model strain *Bacillus subtilis* sp. 3610, which has a well documented specialized metabolome, with the following set of matrices in positive and negative mode: 1,5-diaminonaphthalene (1,5-DAN), 9-aminoacridine (9-AA), ɑ-cyano-4-hydroxycinnamic acid (CHCA), 2,5-dihydrobenzoic acid (DHB), 1:1 CHCA:DHB, 5-formylsalicylic acid (FSA), and 2,4,6-trihydroxyacetophenone (THAP). We and others have had success in using 1,5-DAN matrix for negative mode analyses that target carboxylic acid-containing lipids and metabolites.^19–23^ Likewise THAP has seen success with nucleic acids and nucleotide containing NPs.^24^ Intriguingly, while both CHCA and DHB are common positive mode matrices used for peptides and metabolites in positive mode,^25–28^ we have noted that this same mixture can be utilized in negative mode and significantly increased ionization of cyclic diguanylate monophosphate (c-di-GMP),^29^ a key secondary messenger in a number of microorganisms, demonstrating the need for further exploration for matrices that can facilitate ionization in both positive and negative mode. Finally, while we have observed CHCA:DHB to ionize phenazines and small nitrogen-containing molecules,^29,30^ we also assessed FSA, which is noted for its specificity to ionize alkaloids.^31^

Three MALDI-MS^1^ spectra were acquired with each matrix on a single colony of *B. subtilis sp.* 3610, summed over 3 bursts of 500 laser shots. Following preprocessing of the data and exporting of mass lists (see **Experimental Section**), plotting the average number of detected *m/z* values within replicate spectra across each matrix revealed that the 1:1 CHCA:DHB matrix exhibited the highest number of observed *m/z* values for positive mode analysis, prompting us to move forward with this matrix mixture for positive mode analyses. 1,5-DAN exhibited the highest number of observed *m/z* values for negative mode analysis (**Figure 2A**).^32^

**Figure 2.**
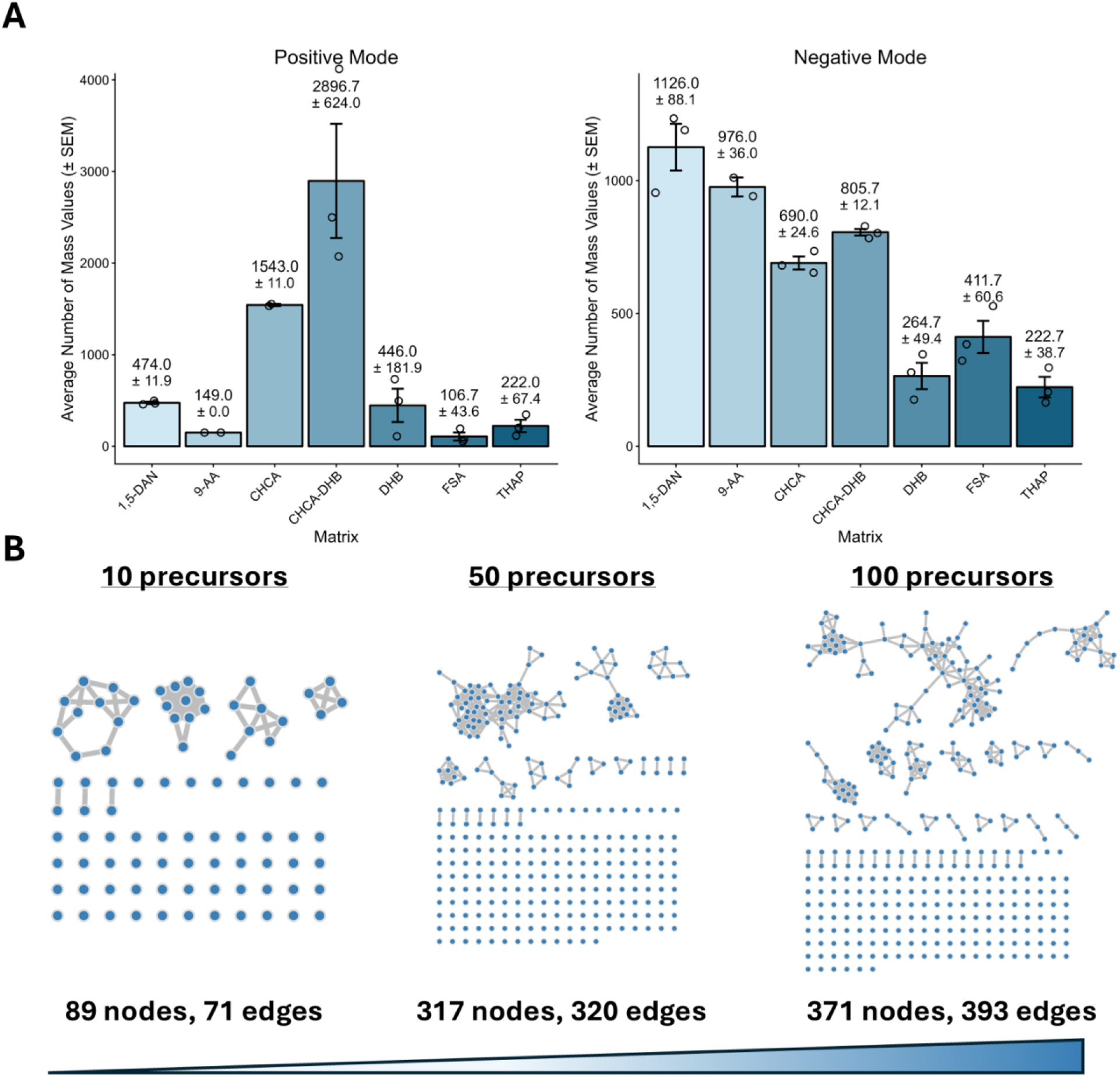
Optimization of MALDI-DDA sample preparation and data acquisition parameters. **A)** MALDI matrix optimization. Bar charts represent the average number of *m/z* features detected across three replicate colonies of *B. subtilis* sp. 2310. Grubbs’ test was applied independently to each matrix group to identify statistically significant outliers (p < 0.05). Replicates identified as outliers were removed before calculation of summary statistics and data visualization. A maximum of one outlier per triplicate set was removed. Error bars represent standard error of the mean (SEM) **B)** A visual representation of the increase in molecular network size as a result of increasing the number of precursors to select for fragmentation via MALDI-MS/MS. The visualized networks are also composed of MALDI-MS/MS data collected from *B. subtilis* colonies.

Initial attempts to generate molecular networks from MALDI-MS/MS data on test samples produced very small networks (89 nodes, 71 edges; **Figure 2B**), largely due to suboptimal precursor selection from the collected MALDI-MS^1^ data. In LC-MS/MS experiments using a QqTOF mass analyzer, it is common to select the top 10 peaks from each MS^1^ scan for fragmentation in subsequent MS/MS scans (i.e., data-dependent acquisition, DDA). Because this process is repeated continuously throughout the chromatographic run, thousands of MS/MS spectra can be collected in a single LC-MS/MS experiment. In contrast, MALDI-MS/MS lacks chromatographic separation and continuous spectral acquisition; instead, a single mass spectrum represents the entire microbial sample. Consequently, selecting only ten precursors results in fragmentation of just ten *m/z* values per sample, sampling only a small fraction of the available chemical space. Increasing the number of selected precursors to 100 *m/z* values substantially expanded network size, enabling more robust molecular networking analysis directly from microbial colonies (**Figure 2**) without sacrificing data quality due to loss of material via repeated laser ablation. We found that beyond the selection of ∼200 *m/z* values, the resulting MS/MS spectra either lacked signal entirely or had poor signal-to-noise ratios. Therefore all subsequent data was collected using 100-200 precursors. Adjusting the number of precursors, alongside the number of laser bursts and shots directly affects the speed of analysis, as each precursor requires its own MALDI laser burst. These parameters should be optimized depending on the user’s needs.

### Targeted Analysis of Marine Actinomycetes

With MALDI-DDA enabled, we first wanted to test the IDBac workflow on a small proof-of-principle subset of bacteria including the marine *Streptomyces* strains sp. CNZ-289 and CNQ-525, known producers of napyradiomycins. Napyradiomycins are halogenated meroterpenoid natural products produced by the *nap* biosynthetic gene cluster (BGC), which encodes the vanadium-dependent haloperoxidases (VHPOs) NapH1, NapH3, and NapH4 that catalyze key halogenation, rearrangement, and cyclization steps.^33–36^ Four additional non-*Streptomyces* Actinomycetota predicted to harbor VHPO-containing BGCs were included in this study: *Actinopolyspora mzabensis* DSM 45460, *Asanoa iriomotensis* DSM 44745, *Asanoa ferruginea* ATCC 49966*, Actinomadura catellatispora* DSM 44772. *Streptomyces* sp. CNZ-289 is also a prolific producer of lavanducyanin, an anticancer phenazinone natural product, which we also sought to monitor in this analysis.^37,38^ Lastly, *B. subtilis* sp. 3610 was included as a “seed” strain for our MALDI-MS/MS analysis, as its metabolite profile is well catalogued.^4,39,40^

Protein MALDI-MS1 (2-20 kDa) and small-molecule MALDI-MS(/MS) (100-2000 Da) were performed directly on colonies of the seven aforementioned strains (see **Experimental Section**). All data were deposited into the IDBac platform for data visualization and analysis. For these and subsequent analyses the protein dendrogram displays cosine distance; in this case, closer to 0 would indicate similarity, whereas large numbers that approach 1 would indicate dissimilarity. Cutoff ranges to display true taxonomic relationships are being actively explored.^13^ The generated protein spectral dendrogram displayed both *Streptomyces* species (CNZ-289 and CNQ-525) as being the most closely related among the deposited strains, albeit with moderate similarity (cosine distance = 0.54; **Figure 3A**). Importantly, our seed strain *B. subtilis sp.* 3610 matched with itself in the IDBac KB (cosine distance = 0.03), indicating that IDBac protein spectral matching functioned as intended. No other strains deposited for this analysis were observed to cluster in any distinct way based on protein spectral similarity.

**Figure 3.**
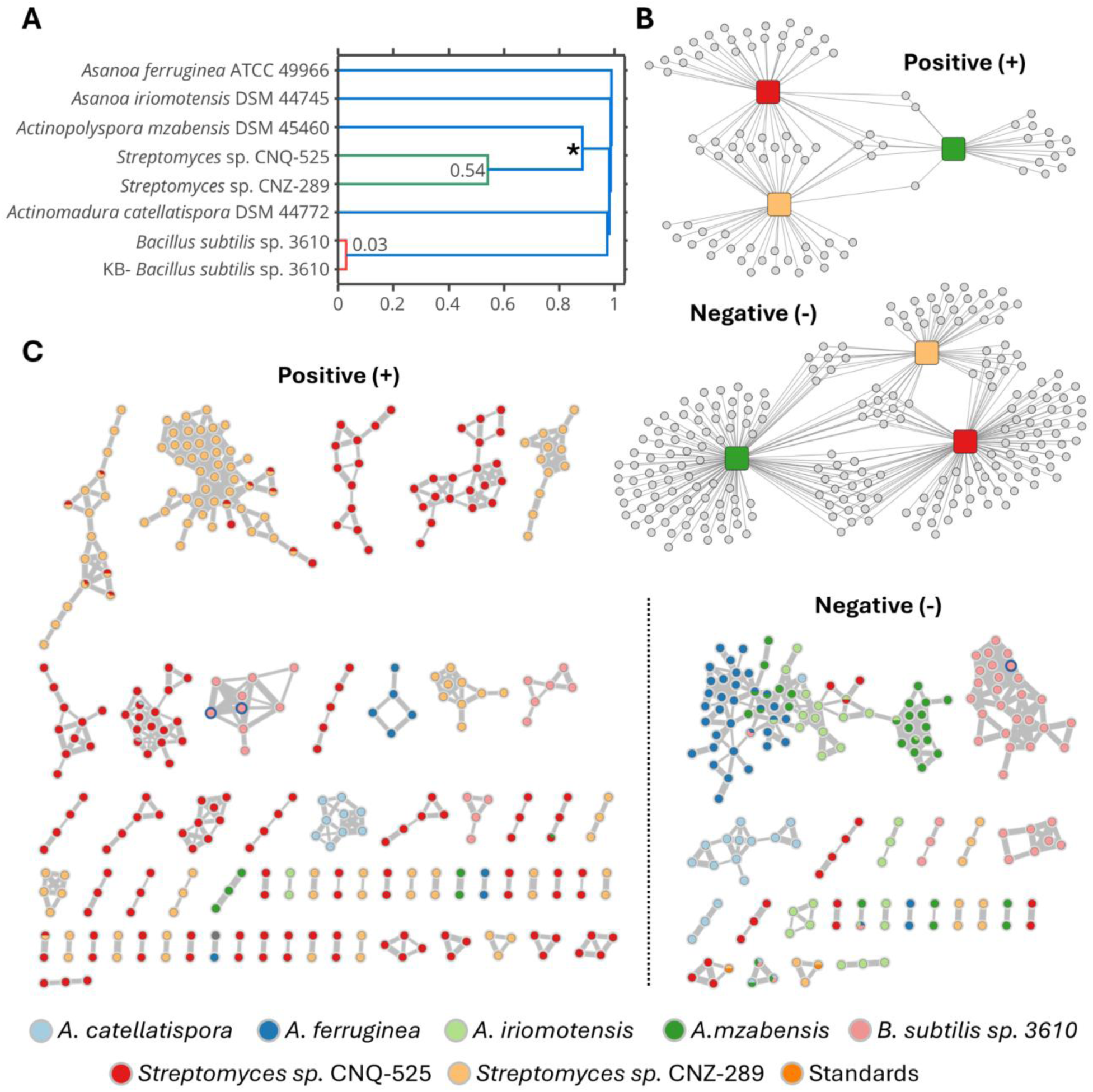
Data output from MALDI-MS IDBac colony analysis. A) Protein spectral dendrogram of the seven strains submitted for analysis. The starred clade represents the cutoff for strains displayed in the positive and negative MANS in panel B. Note that a cosine distance of 0.00 denotes a perfect spectral match and 1.00 represents no match. B) Positive and negative mode MANs for the six actinomycetes + *B. subtilis sp.* 3610. C) Positive and negative mode MS/MS molecular networks generated from MALDI-MS/MS data collected from colonies of the seven strains. Nodes with blue borders indicate GNPS2 library hits.

The positive mode MAN generated from the clade containing both *Streptomyces species* and *A. mzabensis* indicates that *Streptomyces* CNZ-289 and CNQ-525 share more features with each other (25 features) than either share with *A. mzabensis* (7 and 4 features, respectively). Conversely, assessing the same clade in negative mode shifts feature distribution, with *Streptomyces sp.* CNQ-525 sharing more features with *A. mzabensis* (22 features) than with *Streptomyces sp.* CNZ-289 (10 features, **Figure 3B**). This shift is likely the result of the presence of bacterial membrane lipids, which ionize readily in negative ion mode.^41–43^ Despite this, it is likely that some metabolites of interest also ionize efficiently in negative ion mode, making it beneficial to include in the IDBac analysis workflow particularly when using metabolite composition as a metric for constructing diverse microbial libraries. Furthermore, the inclusion of MS/MS data can help mitigate uncertainty regarding the classification of detected metabolites.

To assess whether annotations could be made from the metabolite profiles visualized via MANs, we generated MS/MS precursor lists from the MS^1^ data, followed by MALDI-MS/MS data acquisition. Extending IDBac to include MALDI-MS/MS data upon file import allowed for downstream molecular networking directly from the interactive IDBac web platform on GNPS2. The resulting positive mode molecular network consisted of 312 nodes and 448 edges, while the negative mode network consisted of 170 nodes and 329 edges, both excluding singletons (**Figure 3C, D**). Excitingly, all data presented in **Figure 3** were obtained directly from microbial colonies, with sample preparation, data acquisition, and analysis completed within a single workday (∼8 hours). This emphasizes the extent of dereplication that can be rapidly achieved from single colonies on agar plates.

Additionally, MALDI-MS/MS spectra of purified standards (lavanducyanin, napyradiomycin A1, B1, and B3) were incorporated into the molecular networks as references for these compounds and their analogs. In the positive mode network, a node representing *m/z* 333.206 contained MS/MS spectra from *Streptomyces* sp. CNZ-289 and CNQ 525 along with the lavanducyanin standard (**Figure 4A**). Further examination of the raw MS^1^ data from CNZ-289 and CNQ-525 revealed an ion at *m/z* 333.1961, consistent with the calculated [M+H]⁺ of lavanducyanin (333.1961 Da; mass error = 0 ppm), corroborating the corresponding node observed in the positive-mode MALDI-MS/MS molecular network (**Figure S1**). In the negative mode network, two nodes were observed corresponding to the [M-H]⁻ adducts of napyradiomycin A1 (observed *m/z* 479.138) and B3 (observed *m/z* 557.048), each composed of MS/MS spectra from CNZ-289 and their respective standards (**Figure 4B**). Examination of the negative-mode MS^1^ spectrum of *Streptomyces* sp. CNZ-289 further confirmed the presence of ions at *m/z* 479.1340 (12 ppm error) and 557.0443 (11 ppm error), which exhibited the characteristic dichloro and dichlorobromo isotope patterns consistent with napyradiomycins A1 and B3, respectively (**Figure S2**). Within the generated positive and negative mode networks, surfactins B and C from *B. subtilis sp.* 3610 were annotated from the GNPS2 spectral library, highlighting the ability to use this workflow without verified standards, provided the analytes of interest are included within the spectral library being queried (**Figure 4C**). Lastly, scale up and extraction of *Streptomyces* sp. CNZ-289 led to the isolation of an unreported monochlorinated and dibrominated napyradiomycin B analog (napyradiomycin B8, **1**), the planar structure of which was deduced by MALDI-MS/MS alongside 1D and 2D NMR analyses (**Figures 4D, S3-4, S12-15 Table S5**). B-type napyradiomycins are characterized by a 1,3,6,8-tetrahydroxynaphthalene (THN) core bearing both cyclized C-2 prenyl and C-3 geranyl substituents.^44,45^ The diversity of halogenation across B-type napyradiomycin analogs is well documented, with napyradiomycins B1-7 primarily differentiated by the number and/or position of various halogen and hydroxy substituents.^44–47^ We add to this diversity via the discovery of **1**. Ultimately, this highlights that chemical constituents identified at the colony level can inform the context of isolateable entities upon scale up and extraction.

**Figure 4.**
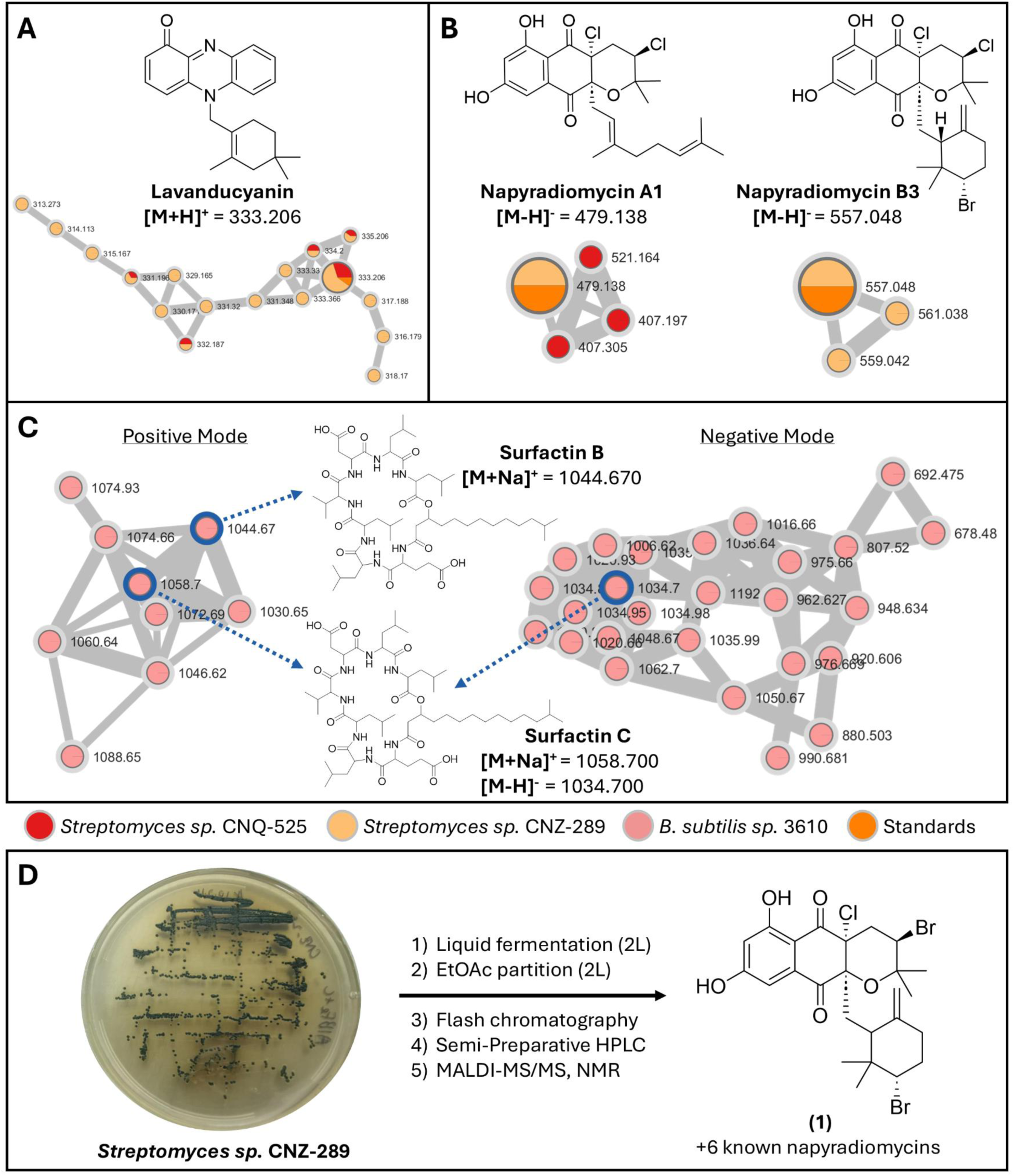
MALDI-MS/MS molecular network annotations from seed standards and GNPS library search. A) Positive mode lavanducyanin molecular family. B) Negative mode napyradiomycin A1 and B3 molecular families. C) Positive and negative mode surfactin molecular families. D) Structure of a newly isolated napyradiomycin B analog (napyradiomycin B8, **1**). We note that the molecular network shows the consensus values for the node and not the precise masses of the ions from individual spectra shown in the supporting information. Stereochemistry is only shown for molecules that have authentic standards, otherwise the stereochemistry is not displayed.

### Untargeted Analysis of Marine Commensal Bacteria

Next the MALDI-MS/MS-enhanced IDBac workflow was tested for its untargeted strain-prioritization capabilities. To do this, 25 commensal bacteria strains that were previously collected from the intestinal contents of various marine fish and mammals in the Monterey Bay were revived from frozen stock and replated for IDBac colony analysis.^14,15^ After cultivation, the colonies were transferred to MALDI target plates and MALDI-MS and MS/MS analysis were performed in a similar manner as described previously. Since the strains used in this analysis were previously identified via 16S rRNA gene sequencing, the generated protein dendrogram was primarily used as additional reference, but pseudotaxonomic clustering was consistent between the 16S rRNA taxonomy and with IDBac KB hits (**Figure S5**). Positive and negative mode MANS allowed for visualization of metabolite association between the 25 strains, and the node counts were considered for strain prioritization as discussed below (**Table 1, Figure S6)**.

**Table 1.** MALDI-MS and MALDI-MS/MS node summary for both positive and negative mode analyses. Strains are listed from top to bottom in order of the highest to lowest number of total unique features from both MS^1^ and MS/MS analysis. *Indicates strains that were prioritized for scale up fermentation, extraction, and fractionation.

| **Taxonomy** | **Strain Code** | **# Unique MS1 Features** | **# Unique MS2 Nodes** |
| --- | --- | --- | --- |
| *Bacillus atrophaeus* | MMA1014 | 12 | 106 |
| *Streptomyces coelicolor* * | MMA1009 | 27 | 78 |
| *Psychrobacter sp.* * | FI1016 | 10 | 40 |
| *Psychrobacter sp.* | FI1008 | 3 | 44 |
| *Rhodococcus luteus* | FI1022 | 16 | 19 |
| *Micromonospora sp.* * | FI1023 | 9 | 26 |
| *Bacillus atrophaeus* | MMA1013 | 8 | 22 |
| *Fictibacillus phosphorivorans* | MMA1027 | 9 | 18 |
| *Paenibacillus sp.* | FI1011 | 6 | 17 |
| *Micrococcinaea bacterium* | FI1004 | 1 | 22 |
| *Bacillus megaterium* | MMA1020 | 5 | 18 |
| *Bacillus firmus* | FI1012 | 12 | 9 |
| *Rhodococcus luteus sp.* | FI1025 | 14 | 5 |
| *Rhodococcus sp.* | FI1018 | 0 | 16 |
| *Arthrobacter sp.* | FI1028 | 9 | 7 |
| *Bacillus oceanisediminis* | MMA1025 | 11 | 4 |
| *Rhodococcus sp.* | FI1015 | 4 | 11 |
| *Bacillus hwajinpoensis* | MMA1010 | 4 | 9 |
| *Rhodococcus luteus* | FI1030 | 5 | 7 |
| *Bacillus oceanisediminis* | MMA1011 | 5 | 7 |
| *Paenibacillus sp.* | FI1009 | 0 | 10 |
| *Micromonospora auratinigra* | MMA1037 | 2 | 6 |
| *Micrococcinaea bacterium* | FI1002 | 1 | 7 |
| *Rhodococcus luteus* | FI1024 | 1 | 6 |
| *Bacillus algicola* | MMA1001 | 3 | 2 |

The combined positive and negative mode MALDI-MS/MS molecular network generated from all 25 strains contained 1420 nodes and 2252 edges, excluding singletons (**Figure S7**). Thirty-nine metabolites were annotated within this network using the GNPS2 library, representing ∼3% of the total nodes within the network. The largest proportion of annotations corresponded to nucleotides (51%) and lipids (23%), which highlights that inclusion of negative mode MS/MS captures lipids that are abundant in the cell membranes; however, several antibiotics were also annotated, including surfactins B and C from various *Bacillus* spp., alongside surugamide A and antimycin A1 from *Streptomyces coelicolor* sp. MMA1009. We suspect the low number of annotations may be attributable to fragmentation differences associated with ionization modality (i.e., differential fragmentation between MALDI- and ESI-MS/MS spectra), which could impact library annotation results.

To demonstrate the prioritization potential of this workflow from colony-based materials, we used the MALDI MANs and molecular networks for the 25 strains to create a ranked prioritization based on taxonomic representation and the number of strain-specific features at both the MS^1^ and MS/MS levels for each strain (i.e., nodes not shared with any other strain in the respective network) (**Table 1**). We posit that this approach could enable strains to be prioritized not only by the number of unique nominal *m/z* values detected, but also by the richness of potentially unique molecular entities through the inclusion of MS/MS analysis.

After assessing strains based on their unique *m/z* features, we prioritized three strains for further scale-up, fractionation, and LC-MS/MS analysis based on taxonomic considerations: *Streptomyces coelicolor* sp. MMA1009 for its well-characterized metabolome and underrepresentation of *Streptomyces* within our dataset (serving as a pseudo-control for metabolite annotation), *Psychrobacter* sp. FI1016 due to the limited representation of *Psychrobacter*-associated metabolites in common NP databases (e.g. NP Atlas^48^, Dictionary of Natural Products, and COCONUT^49^) and *Micromonospora* sp. FI1023 due to the limited representation of the genus within the strain collection used during this analysis. Despite possessing the highest total number of metabolite features, *Bacillus atrophaeus* sp. MMA1014 was excluded from prioritization due to the high representation of *Bacillus* species within our dataset. At this stage, strain prioritization using the IDBac x GNPS2 platform would be considered complete based on the low input biomass of a single colony, and subsequent fermentation, fractionation and downstream analyses would be left to the discretion of the researcher. It is important to reiterate that subsequent studies are required, and this workflow is both complementary and orthogonal to other techniques and approaches like NMR, X-ray crystallography, microED, and bioactivity-guided fractionation.

In the case of this study, the prioritized trains were re-cultivated and scaled up to 1L cultures and subjected to XAD-16N resin extraction. The resulting crude extracts were further fractionated into 7 subfractions using solid phase extraction (SPE). For each strain, all subfractions were subjected to MALDI-MS/MS (via MALDI-DDA) and LC-MS/MS analysis in positive ion mode and the resulting datasets were combined into molecular networks constructed from both MALDI- and LC-MS/MS data for each prioritized strain. From the combined networks, *Streptomyces coelicolor* sp. MMA1009 displayed 213 library annotations, *Psychrobacter* sp. FI1016 displayed 183 library annotations, and *Micromonospora* sp. FI1023 displayed 150 library annotations. Of the three strains, *Streptomyces coelicolor* sp. MMA1009 additionally exhibited the highest number of bioactive natural product annotations, including surugamides A and G (anticancer^50^, antifungal^51^, and antimicrobial^52^), desferrioxamine and dehydroxynocardamine (siderophores^53,54^), albubactin acid A (anticancer^55^) and antimycin A analogs (antifungal, insecticidal, and nematocidal^56^)(**Figure 5A, S8-9**).

**Figure 5.**
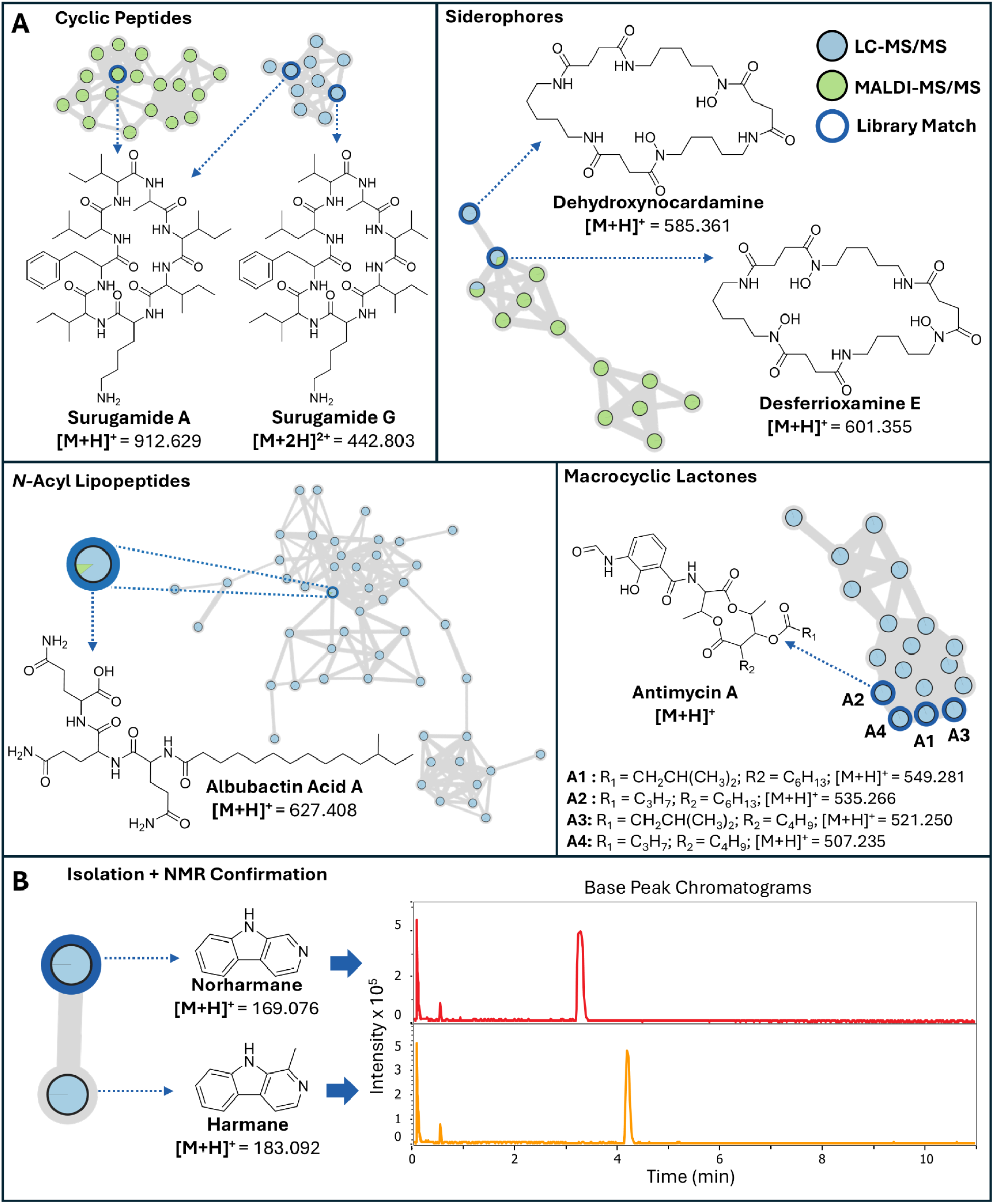
Combined MALDI- and LC-MS/MS molecular network annotation results from SPE fractions of marine commensal bacteria extracts. **A)** GNPS2 molecular families and library hits for small molecules produced by *Streptomyces coelicolor sp.* MMA1009. **B)** GNPS2 molecular family, and subsequent base peak chromatograms collected after the isolation of harmane and norharmane from *Psychrobacter sp.* FI1016.

Notably, several of these annotations were detected by both MALDI- and LC-MS/MS (surugamide A, desferrioxamine E, and albubactin acid A). Despite antimycin A not being detected via MALDI-MS/MS in the co-network analysis, re-detection of both antimycin A and surugamide A corroborate the annotations observed from *Streptomyces coelicolor* sp. MMA1009 at the colony level, further supporting that colony-based MALDI-MS/MS can inform the context of metabolite composition upon scale up and extraction. Finally, mass-guided fractionation of *Psychrobacter* sp. FI1016 led to the successful isolation of the tryptamine-derived neurochemicals β-carboline (norharmane) and 1-methyl-β-carboline (harmane), which were confirmed through LC-MS/MS fragmentation, and 1 and 2D NMR analyses (**Figure 5B**, **S10-11**, **S17-20**, **Tables S6-7**). While these metabolites were not initially observed at the colony level, a mass-guided approach following strain prioritization using only MALDI-MS techniques led to the detection and isolation of bioactive chemical entities.

### Considerations

Limitations to the broader IDBac platform including cultivation conditions, choice of matrix, and specific limitations to IDBac-guided library building have been discussed in detail previously.^4,5,13^ With regard to the collection of MALDI-MS/MS data directly from microbial colonies, a number of mass analyzers could be equipped with atmospheric pressure (AP)-MALDI sources or are readily commercially available. However, the isolation windows will vary based on the mass analyzer specifications and the sensitivity between AP-MALDI and sources under vacuum. Wide isolation windows, such as those on MALDI-TOF/TOF instruments can lead to the fragmentation of unrelated ions, resulting in chimeric fragment peaks that can decrease annotation accuracy when querying spectral libraries. MALDI analysis can be more susceptible to this due to the lack of chromatographic separation prior to ionization. The addition of ion mobility spectrometry (IMS) as a timescale-compatible method for gas-phase ion separation could help reduce chimeric spectra and enhance annotation, but requires further specialized equipment.^57,58^ Due to software limitations and the nature of the MALDI-DDA analysis described herein, fragmentation of analytes can only be performed at a single collision energy (CE) per acquisition. Running replicate plates at different CE values could address this limitation, but sacrifices overall throughput. MzMine^59^ allows the scheduling of MALDI-MS/MS spectra at varying collision energies, but requires the use of trapped ion mobility (TIMS) to perform these specialized workflows.^60,61^ Lastly, when fragmenting NP standards via collision induced dissociation (CID) across multiple collision energies (15-75 eV) with both ESI and MALDI ionization, we observed that in some cases fragmentation spectra appeared to be fundamentally different for the same analyte across ionization modalities (**Figure S21**). This could potentially lead to further loss of annotation abundance and accuracy when querying MS/MS spectral libraries, which are primarily constructed of LC-MS/MS data.

## Conclusions

In this study we showcase the use of MALDI-MS/MS to provide molecular annotation at the colony-level, further enhancing high-throughput microbial dereplication and decision making via the IDBac x GNPS2 platform. In a targeted example, we successfully observed the production of lavanducyanin, napyradiomycin A1, and napyradiomycin B3 from *Streptomyces sp.* CNZ-289 using MALDI-MS/MS and downstream molecular networking from IDBac. Further investigations of a large-scale extract of *Streptomyces sp.* CNZ-289 led to the isolation of an unreported monochlorinated and dibrominated napyradiomycin B analog (napyradiomycin B8, **1**). Extending this workflow to an untargeted investigation of 25 marine commensal bacteria strains led to the prioritization of 3 strains for further analysis within a single workday. This ultimately resulted in the re-detection of surugamide A, and antimycin A from *Streptomyces coelicolor* sp. MMA1009, corroborating initial colony-based annotations and the isolation of harmane and norharmane from a marine *Psychrobacter sp.*. Together, these results suggest that MALDI-MS/MS directly from microbial colonies can enhance the decision-making process prior to scale up, extraction, and fractionation.

## Experimental Section

### Materials

*Streptomyces* spp. CNZ-289 and CNQ-525 were gifted by Paul Jensen and Doug Sweeney at the Scripps Institution of Oceanography (La Jolla, CA). *Actinopolyspora mzabensis* (DSM 45460), *Asanoa iriomotensis* (DSM 44745), and *Actinomadura catellatispora* (DSM 44772) were purchased from DSMZ (Braunschweig, Germany). *Asanoa ferruginea* (ATCC 49966) was purchased from ATCC (Manassas, VA, USA). *Bacillus subtilis sp.* 3610 was obtained from Pieter Dorrestein at UCSD. Purified standards of napyradiomycins A1 and B3 were already in our in-house library. Protein Calibration Standard 1 was purchased from Bruker Daltonics (Billerica, MA, USA). Phosphorus red (CAS: 7723-14-0) MALDI-MS calibrant was purchased from Spectrum Chemical Mfg. Corp. (Gardena, CA, USA).

### Bacterial Strain Cultivation

All strains were removed from a −70 °C freezer and a small amount of each frozen stock was transferred to an agar plate and streaked using sterile inoculation loops. The marine *Actinomycetota* from our targeted analysis were cultivated on A1BFe+C agar (**Table S2**) and incubated inverted at 30 °C for 10-14 days until colony formation was observed. The 25 marine commensal bacteria strains and *B. subtilis* were cultivated on Difco Marine Agar 2216 and incubated inverted at 24 °C for 7 days until colony formation was observed.^14,15^

### MALDI Matrix Optimization

Solutions of 1,5-DAN, 9-AA, CHCA, DHB, 1:1 CHCA:DHB, FSA, and THAP were prepared (50 mM in 78:22 Acetonitrile:MilliQ H2O + 0.1% TFA). To test each matrix, three colonies of *B. subtilis* were spread onto a MALDI target plate per matrix. Each matrix was then spotted (1 µL per colony spot) on top of each respective triplicate group of *B. subtilis* colonies. MALDI spectra were collected as outlined in **MALDI-MS and MS/MS Acquisition** below. For each spectrum, a mass list was generated in Bruker’s DataAnalysis software using the MassList “Find” function with the signal-to-noise threshold set to 4, the relative intensity threshold (base peak) set to 0.1%, and the absolute intensity threshold set to 100. The number of detected *m/z* features for each matrix was then averaged over the three replicates and the averages plotted in bar charts as shown in **Figure 2A**. To assess replicate consistency, Grubbs’ test was applied independently to each matrix group to identify statistically significant outliers (p < 0.05). Replicates identified as outliers were removed before calculation of summary statistics and data visualization. A maximum of one outlier per triplicate set was removed.

### MALDI Protein MS Sample Preparation

Five colonies of each strain were transferred to individual spots in five-spot groups on a MALDI target plate using autoclaved toothpicks. Aqueous formic acid (1 µL of 70%) was spotted atop each colony and the MALDI target was allowed to dry in a chemical fume hood. Then, 1 µL of 50 mM sinapic acid was spotted atop each colony. For each five-spot group, 1 µL of a 1:1 mixture of Protein Calibration Standard 1 and 50 mM sinapic acid was spotted as the protein-MS calibrant and the MALDI target was once again allowed to dry in a chemical fume hood.

### MALDI Small Molecule MS Sample Preparation

Five colonies for each strain were prepared for MALDI small molecule MS and MS/MS analysis by forgoing the application of 70% formic acid, and directly applying 1 µL of 50 mM 1:1 α-cyano-4-hydroxycinnamic acid:2,5-dihydrobenzoic acid (CHCA:DHB) for positive mode analysis, or 1 µL of 50mM 1,5-diaminonaphthalene (1,5-DAN) for negative mode analysis. Lastly, 1 µL of saturated phosphorous red was spotted on the MALDI target plate for MS calibration.

### MALDI-MS and MS/MS Acquisition

Protein MS data were collected on a MicrofleX LT (Bruker Daltonics) linear MALDI-TOF mass spectrometer in positive ion mode from 2-20 kDa. MALDI ionization on the MicrofleX was performed with an N2 laser operating at 337 nm and a frequency of 60Hz. Shots were summed in 200 shot intervals until 600 shots above intensity 200 were acquired. Small molecule MS data were collected on a timsTOF fleX QqTOF mass spectrometer (Bruker Daltonics) in positive and negative ion modes from 100-2000 Da. The laser was set to a frequency of 5000 Hz, and data acquired in 5 bursts of 200 shots (total of 1000 shots per spot). Laser pitch was set to 100 x 100 µm to maximize sampling area, and the MALDI stage movement was set to “random walk” to allow for randomized sampling across each sample spot. For the collection of MS/MS data, the mass range was expanded to 50-2000 Da to account for fragmentation of low molecular weight species (100-200 Da) selected by the python script for fragmentation. CEs and isolation windows were interpolated on the basis of *m/z*, ranging from a CE of 20.00 eV and an isolation width of 1.00 *m/z* at 50 *m/z* to a CE of 70.00 eV and an isolation width of 8.00 *m/z* at *m/z* 2000. All other parameters remained the same between MS^1^ and MS^2^ analysis.

### Data conversion and IDBac Analysis

All MALDI- and LC-MS data were converted into .mzML files using MSConvert from Proteowizard^62^ with vendor peak picking enabled for the respective MS^n^ level to ensure peak centroiding for small molecule data. Protein MS data were not centroided during data conversion, as centroiding occurs during the IDBac protein MS data processing. MALDI protein small molecule data were uploaded into the File Browser on GNPS2 via drag and drop. The protein/small molecule metadata table was filled out as outlined in the IDBac analysis documentation.^13^ Protein data were processed using the “idbac_split_maldi_workflow” to organize spectra from biological replicates into separate .mzML files. For small molecule data, associated MS/MS files were merged with their respective MS^1^ .mzML files using a newly developed interactive interface (see Supplementary Materials and Methods). To include MS/MS data into the IDBac analysis, the “smol_mol_contains_ms2” parameter was set to “yes”. IDBac interactive protein analysis was performed for all datasets using a mass range of 2000-20000 Da (2-20kDa) and a 10 Da bin size. The distance metric was set to “cosine”, with the database search cosine threshold set to 0.5. For the construction of MANS, the relative intensity threshold was set to 0.02; the replicate frequency threshold was set to 0.6 (i.e. an *m/z* value must be present in 3 out of 5 replicates to be included in the MAN); and the mass range was set to 100-2000 *m/z* with a tolerance of 0.05 *m/z*.

### Downstream Molecular Networking

From the IDBac interactive analysis task page, “Downstream Analysis - Molecular Networking” was selected. From there, classical molecular networking was performed as normal in GNPS2. The data was filtered by removing all MS/MS fragment ions within +/- 17 Da of the precursor *m/z*. MS/MS spectra were window filtered by choosing only the top 6 fragment ions in the +/- 50Da window throughout the spectrum. The precursor ion mass tolerance was set to 0.02 Da and a MS/MS fragment ion tolerance of 0.02 Da. A network was then created where edges were filtered to have a cosine score above 0.70 and more than 6 matched peaks. Further, edges between two nodes were kept in the network if and only if each of the nodes appeared in each other’s respective top 10 most similar nodes. Finally, the maximum size of a molecular family was set to 100, and the lowest scoring edges were removed from molecular families until the molecular family size was below this threshold. The spectra in the network were then searched against GNPS2’s spectral libraries. The library spectra were filtered in the same manner as the input data. All matches kept between network spectra and library spectra were required to have a score above 0.70 and at least 6 matched peaks. The molecular networks were exported as .graphml files and visualized Cytoscape (v 3.10.1). Nodes were color coded by strain as indicated, and edge width was scaled to cosine score. Nodes containing spectra from media fractions were removed from the networks by applying a column filter and deleting the selected nodes.

### Large-Scale Fermentation, Extraction, and Fractionation

For *Streptomyces* sp. CNZ-289, spores from the colony were inoculated into 100 mL of A1BFe+C liquid medium and cultivated at 30°C for 5 days. The seed culture (80 mL) was transferred to 2 L of A1BFe+C liquid medium and cultivated at 30 °C for 10 days. Cultures were incubated with glass beads and shaken at 200 rpm. The 2 L culture was then partitioned with 4 L of EtOAc, and the organic layer was evaporated to dryness on a rotary evaporator to obtain a crude extract. The extract was subjected to fractionation with silica gel flash chromatography (hexane/EtOAc/MeOH, gradient 10:1:0 to 0:0:1) resulting in 10 fractions designated as A-J based on increasing polarity of eluent. The resulting fractions were evaporated to dryness using a rotary evaporator.

For the three prioritized commensal bacteria (MMA1009, FI1016, FI1023), selected colonies of each strain were inoculated into 10 mL of Difco Marine Broth 2216. The 3 cultures were stepped up in stages at 7-day intervals by first inoculating 1.5 mL of the 10-mL cell cultures into 50 mL Difco marine broth (medium scale), followed by inoculation of 40 mL of these medium-scale cell cultures into 1 L of the same media broth, now containing 20.0 g Amberlite XAD-16 adsorbent resin in 2.8 L Fernbach flasks for 7 days. All cultures were incubated at 25 °C containing glass beads and shaken at 200 rpm. This procedure was performed in duplicate for *Streptomyces coelicolor* sp. MMA1009 with ISP2 broth (**Table S1**). The cells and resin were collected from the bacterial extract by vacuum filtration using Whatman glass microfiber filters and washed with deionized water. This cell/resin slurry was extracted with 250 mL of 1:1 MeOH/DCM, and the organic extract was removed by vacuum filtration and concentrated to dryness using a rotary evaporator. The crude organic extracts were subjected to solid-phase extraction using a SupelcoDiscovery C18 cartridge (10 g) and eluted using a step gradient of 80 mL MeOH/H2O solvent mixtures (10% MeOH (A), 20% MeOH (B), 40% MeOH (C), 60% MeOH (D), 80% MeOH (E), 100% MeOH (F), and ethyl acetate (G)) to afford seven fractions designated as A−G. The resulting fractions were evaporated to dryness using a rotary evaporator.

### HPLC Fractionation

Flash chromatography fractions A and B (hexane/EtOAc, gradient 10:1 - 10:4) of *Streptomyces* sp. CNZ-289 were subjected to semi-preparative reverse-phase HPLC (Phenomonex Luna 5 µm C18(2) 100 Å, 250 x 10 mm, 85:15 ACN:H2O isocratic run over 16 min, 2.00 mL/min flowrate) to produce the napyradiomycin B8 analog, which eluted at 15.7 min.

Fraction E from *Psychrobacter* sp. Fish1008 was subjected to semi-preparative reverse-phase HPLC (Phenomonex Kinetix 5 µm C18 100 Å, 250 x 10mm, gradient run: 10:90 ACN:H2O from 0.00-4.15 min; raise to 25:75 ACN:H2O from 4.15-15.82 min; raise to 30:70 ACN:H2O from 15.82-20.00 min; raise to 100:0 ACN:H2O from 20:00-21.00 min; hold 100:0 ACN:H2O from 21.00-24:00 min; lower to 10:90 ACN:H2O from 24.00-25.00 min; hold 10:90 ACN:H2O from 25.00-30.00 min; 4.73 mL/min flowrate) to yield β-carboline at 11.02 min and 1-methyl-β-carboline at 12.75 min.

## Author Notes

## Acknowledgements

This work was supported by the National Institute of General Medical Sciences of the NIH award R21GM148870 (LMS), R25GM051765 (CDS), R35GM147235 (SMKM), and by National Science Foundation graduate research fellowship program (RAS). MS, MW were supported by NIH 5U24DK133658-02; and the National Institute of Allergy and Infectious Diseases under award number F31AI200270 (MS). MS was supported by the National Science Foundation and the NSF ExFab Biofoundry under award no. DBI-2400327. The content is solely the responsibility of the authors and does not necessarily represent the official views of the National Institutes of Health.

## Data Availability

### Raw Data

All MALDI- and LC-MS data in this study are available under CC0 1.0 Universal License as raw (*.d) and open source (*.mzML) data formats: doi:10.25345/C50863K7S. MassIVE accession: MSV000101998. All NMR data have been deposited to NP-MRD:

### Molecular networks

Marine Actinomycetota (negative mode MALDI): https://gnps2.org/status?task=9a7f038d56294a03bf6413b41c4201f6

Marine Actinomycetota (positive mode MALDI): https://gnps2.org/status?task=3cddcbada232426aa4cb77089cf70909

Marine Commensal Bacteria (positive + negative mode MALDI): https://gnps2.org/status?task=b6adf5d2a2c842ccaec3979f02385f13

MMA1009-ISP2 Combined SPE Fraction Network (positive mode MALDI + LC-MS/MS): https://gnps2.org/status?task=8c5fc0a10cb64cac88372586d8a40a6b

MMA1009-Marine Broth Combined SPE Fraction Network (positive mode MALDI + LC-MS/MS): https://gnps2.org/status?task=41fa329afe164937b60ce329612eb085

FI1016-Marine Broth Combined SPE Fraction Network (positive mode MALDI + LC-MS/MS): https://gnps2.org/status?task=0acc9fbf1f6144f1849c2fb1994dddd6

FI1023-ISP2 Combined SPE Fraction Network (positive mode MALDI + LC-MS/MS): https://gnps2.org/status?task=ef9dc8e8e4f246f587e2252e0420c694

### Scripts and GUI

All tools for MALDI-MS/MS precursor selection on the timsTOF fleX and be found here: https://github.com/gtluu/flex_maldi_dda_automation

The specific tool for MALDI-MS/MS precursor selection can be found here: https://gtluu.github.io/flex_maldi_dda_automation/msms_autox_generator.html

## Supporting Information

- Media recipes, additional molecular networks, and all instrument parameters.

Characterization data, including HR-MS, MS/MS, and 1D and 2D NMR spectra of isolated compounds.

