## Supporting Information for "MALDI Tandem Mass Spectrometry for Colony-Based Dereplication of Natural Products"

**Table S1.** ISP2 Media Recipe…………………………………………………………… S2
**Table S2.** A1BFe+C media Recipe……………………………………………………… S3
**Table S3.** Artificial Seawater Recipe…………………………………………………… S4
**Table S4.** Difco Marine Broth Recipe ………………………………………………… S5
**Software Development**………………………………………………………………… S6
**Figure S1.** MALDI-MS[1] Spectrum of lavanducyanin from *Streptomyces* sp. CNZ-289 colonies.. S7
**Figure S2.** MALDI-MS[1] Spectra of napyradiomycins A and B from *Streptomyces* sp. CNZ-289 colonies……………………………………………………………………… S8
**Figure S3.** MALDI-MS[1] spectrum of napyradiomycin B8……………………………… S9
**Figure S4.** MALDI-MS/MS spectra of compound 1 vs napyradiomycin B3 standard……….... S10
**Figure S5.** Protein spectral dendrogram of 25 commensal microbial strains…………….... S11
**Figure S6.** Positive and negative mode MANS of 25 commensal microbial strains………… S12
**Figure S7.** Combined positive and negative mode MALDI-MS/MS molecular network of 25 commensal microbial strains…………………………………………………… S13
**Figure S8.** Combined MALDI- and LC-MS/MS molecular network of SPE fractions A-F of *Streptomyces coelicolor* sp. MMA1009…………………………………………… S14
**Figure S9.** Mirror plots of combined MALDI- and LC-MS/MS spectra……………………….. S15
**Figure S10.** Combined MALDI- and LC-MS/MS molecular network of SPE fractions A-F of *Psychrobacter sp.* FI1016……………………………………………………….. S16
**Figure S11.** MS/MS spectral comparison of harmane and norharmane…..……………...…….. S17
**Figure S12.** Combined MALDI- and LC-MS/MS molecular network of SPE fractions A-F of *Micromonospora sp.* FI1023……………………………………………………….. S18
**Table S5.** $^{1}$H NMR comparison of the isolated napyradiomycin B8 (**1**) with the most recently reported $^{1}$H NMR data of napyradiomycin B1……………………………….... S20
**NMR Spectra for napyradiomycin B8**…………………………………………………….. S21-24
**Table S6.** $^{1}$H NMR comparison of the isolated norharmane (**2**) with the values reported by Siderhurst et al. in $CD_3OD$…………………………………………………… S26
**NMR Spectra for norharmane (2)**………………………………………………………… S27-28
**Table S7.** $^{1}$H NMR comparison of the isolated harmane (**3**) with the values reported by Seki et al. in $CD_3OD$…………………………………………………………………… S29
**NMR Spectra for harmane (3)**...……………………………………………………………… S30-31
**Figure S21** Comparison of fragmentation of andrastin A between MALDI- and ESI-MS/MS at 15 and 25 eV…………………………………………………………………………... S32
**References**……………………………………………………………………………….. S33

### Materials and Methods

#### Media Recipes

**Table S1.** Recipe for ISP2 media. Liquid media recipe is the same without agar.

| ISP2 Component | Amount (per 1L) |
|---|---|
| Yeast extract | 4.0 g |
| Malt extract | 10.0 g |
| Dextrose | 4.0 g |
| Agar | 15.0 g |
| MilliQ $H_2O$ | 1.0 L |
| pH adjusted to 7.2 using 1.0 M NaOH. Autoclaved at 121°C for 20 minutes | |

**Table S2.** Recipe for A1BFe+C media. Liquid recipe is the same without agar.

| A1BFe+C Component | Amount (per 1L) |
|---|---|
| Soluble starch | 10.0 g |
| Yeast extract | 4.0 g |
| Peptone | 2.0 g |
| Calcium carbonate | 1.0 g |
| KBr | 100 mg |
| $Fe_2(SO_4)_3·4H_2O$ | 40 mg |
| Agar | 15.0 g |
| Seawater | 1L |
| Autoclaved at 121°C for 20 minutes | |

**Table S3.** Artificial seawater recipe.

| Seawater component | Amount (per 1L) |
|---|---|
| NaCl | 21.2 g |
| $Na_2SO_4$ | 3.5 g |
| KCl | 0.6 g |
| $NaHCO_3$ | 0.175 g |
| KBr | 0.1 g |
| Boric acid | 23 mg |
| NaF | 3.0 g |
| $MgCl_2 \cdot 6H_2O$ | 9.6 g |
| $CaCl_2 \cdot 6H_2O$ | 1.3 g |
| $SrCl_2$ | 22 mg |
| $NaNO_3$ | 2.3 mg |
| $NaH_2PO4 \cdot H_2O$ | 0.16 mg |
| $FeCl_3 \cdot 6H_2O$ | 0.09 mg |
| $Na_2EDTA \cdot 2H_2O$ | 0.28 mg |
| $ZnSO_4 \cdot H_2O$ | 3.6 μg |
| $CoSO_4 \cdot 7H_2O$ | 0.5 μg |
| $MnSO_4 \cdot H2O$ | 27 μg |
| $Na_2MoO_4 \cdot 2H_2O$ | 0.07 μg |
| $Na_2SeO_3$ | 0.09 ng |
| $NiCl_2 \cdot 6H_2O$ | 0.075 μg |
| thiamine-HCL | 5 μg |
| biotin | 0.1 μg |
| Vitamin B12 | 0.05 μg |
| Autoclaved at 121°C for 20 minutes | |

**Table S4.** Difco Marine Broth recipe.

| Marine Broth Component | Amount (per 1L) |
|---|---|
| Difco Marine Broth | 37.4 g |
| MilliQ $H_2O$ | 1L |
| Autoclaved at 121°C for 20 minutes | |

### Software Development

#### Graphical User Interface for MALDI-DDA Precursor List Generation

Several automation tools have been developed for the timsTOF fleX platform (https://github.com/gtluu/flex_maldi_dda_automation). Among these tools, a workflow titled “fleX MS/MS AutoXecute Generator” with a graphical user interface (GUI) has been developed to allow for semi-automated precursor selection from MALDI-qTOF dried droplet MS datasets in a data dependent fashion on the Bruker timsTOF fleX. The GUI is a dash and pywebview application that was developed in Python 3.11. Users can load full scan MALDI MS1 datasets acquired via AutoXecute mode in timsControl and process the data as described below. Since this workflow uses local filepaths, it is recommended to run it on the computer where the data was originally acquired and stored. However, if the workflow is run on a different computer, users may be required to manually search for or change filepaths in the GUI.

Prior to precursor selection, the full scan MS1 can then be preprocessed using user specified parameters and previewed. Replicate spectra can also be grouped to generate a consensus spectrum prior to (pre)processing, and control spectra can be used to generate a precursor exclusion list. For more detailed information, please refer to the documentation: https://gtluu.github.io/flex_maldi_dda_automation/msms_autox_generator.html

Once the MS1 data has been processed, a new AutoXecute file will be generated. This file can be loaded into Bruker timsControl, and the Validate button in timsControl’s AutoXecute window can be used to ensure there are no issues with the generated file.

#### Incorporation of MALDI-MS/MS Data into the IDBac web platform

MALDI-MS/MS was directly integrated into the IDBac web platform via the IDBac MS/MS association tool and the IDBac Analysis Workflow. Concretely, using a newly developed interactive interface (https://idbac.org/MSMSFileGenerator), MS/MS files produced by the MALDI-DDA protocol are merged into the small molecule .mzML files to generate pseudo-run files. These files are processed by the IDBac analysis workflow into a .MGF format compatible with GNPS2, facilitating downstream analysis of MALDI-MS/MS data that mimics common DDA LC-MS/MS analysis workflows. Finally, the workflow enables an automated hand-off to Classical Molecular Networking, allowing users to launch the analysis as a pre-configured step without additional manual data entry.

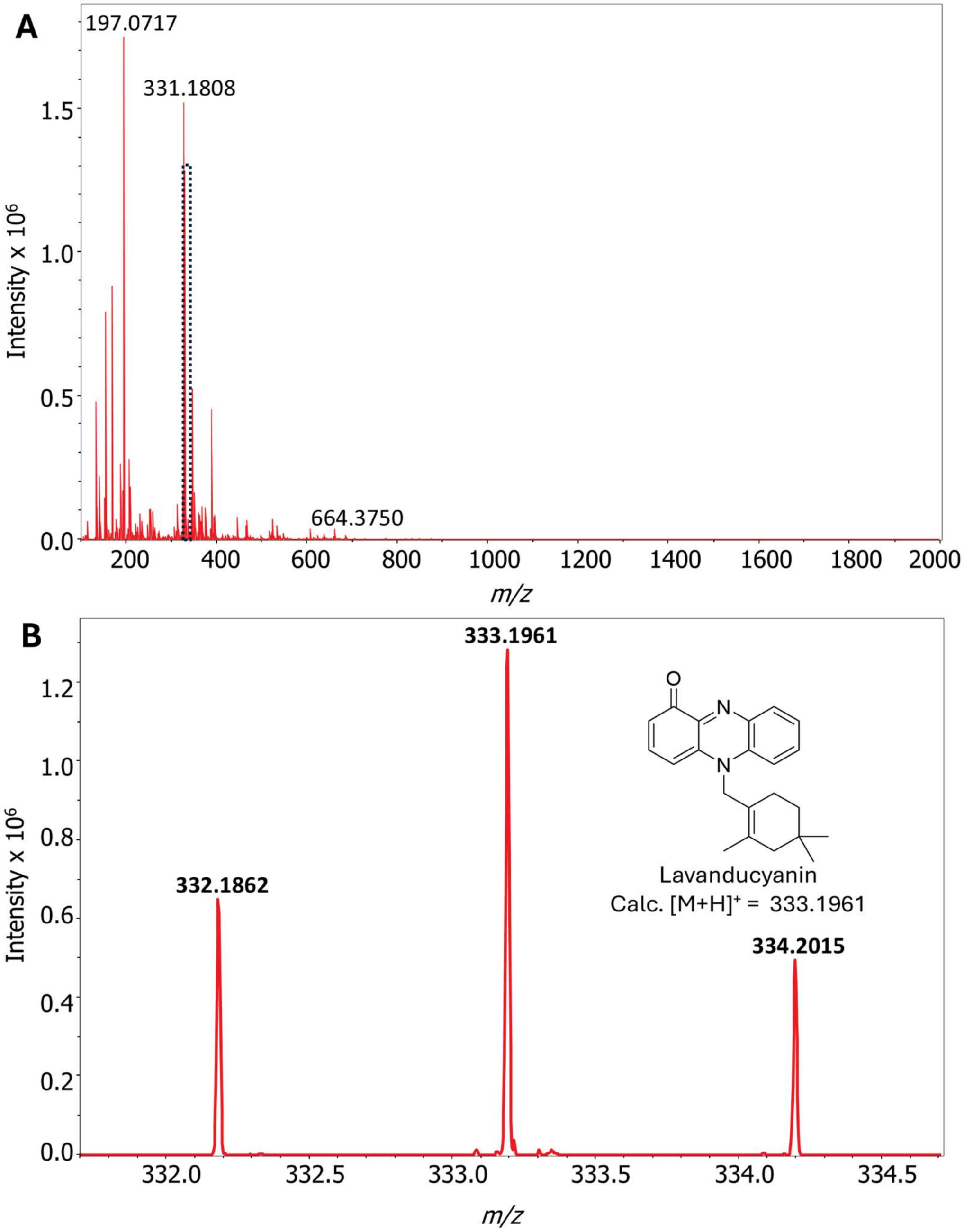


**Figure S1** MALDI-MS[1] spectrum of lavanducyanin collected directly from *Streptomyces sp.* CNZ-289 colonies. **A)** Full scan MALDI-MS[1] spectrum of CNZ-289. **B)** Zoomed in MALDI-MS[1] spectrum of lavanducyanin. Lavanducyanin can be ionized in two different forms, as the protonated molecule $[M+H]^+$ and as the molecular ion $[M]^{\cdot+}$ Both ions are readily observed as well as the $^{13}C$ isotopologue. Consistent with the calculated $[M]^{\cdot+}$ of lavanducyanin (332.1883 Da; mass error = -6.3 ppm)

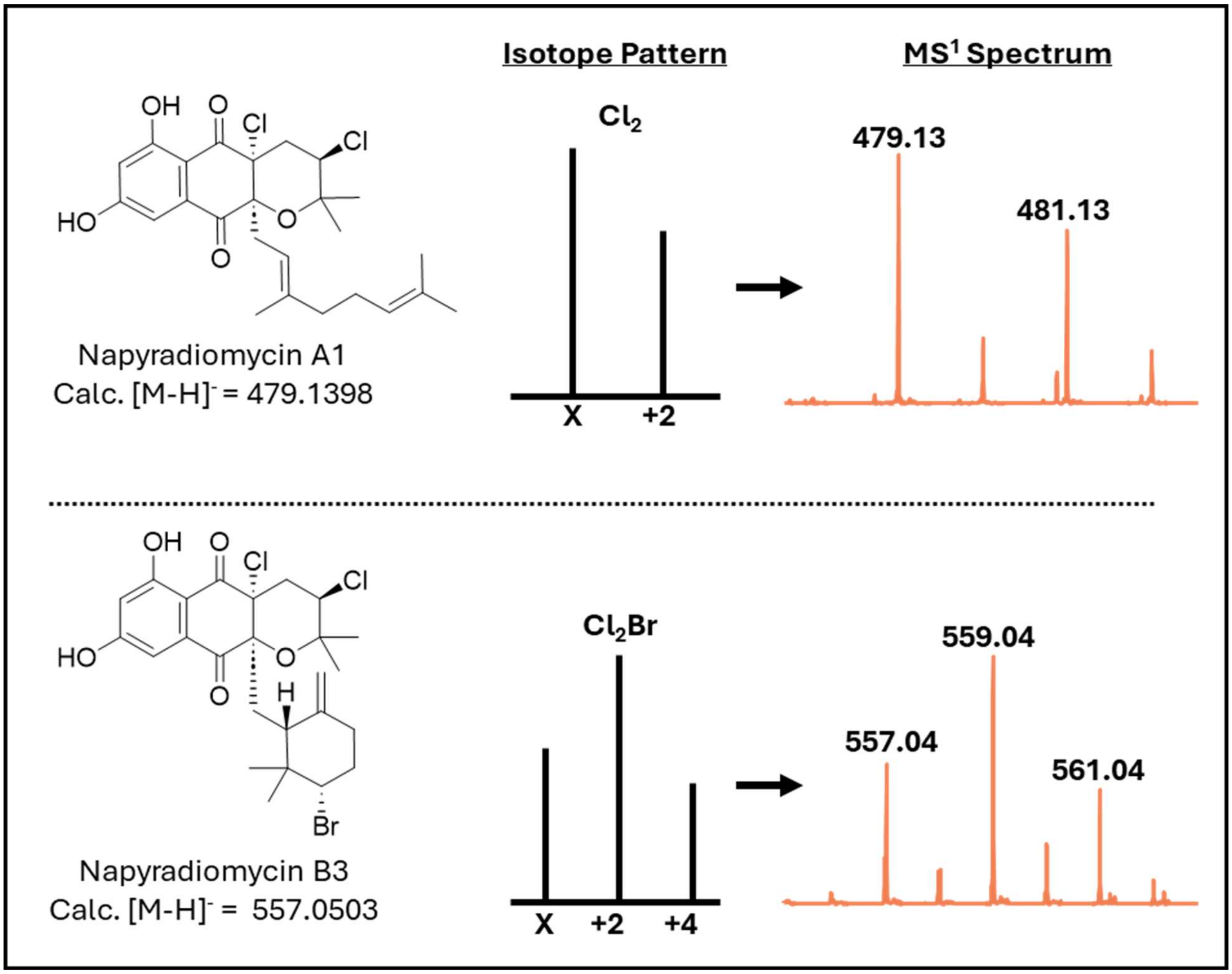


**Figure S2** MALDI-$MS^1$ spectra of napyradiomycins A1 and B3 collected directly from *Streptomyces sp.* CNZ-289 colonies. The diagnostic isotopic distribution for the respective halogenation scheme is shown on the left side, and the experimental spectrum is shown on the right.

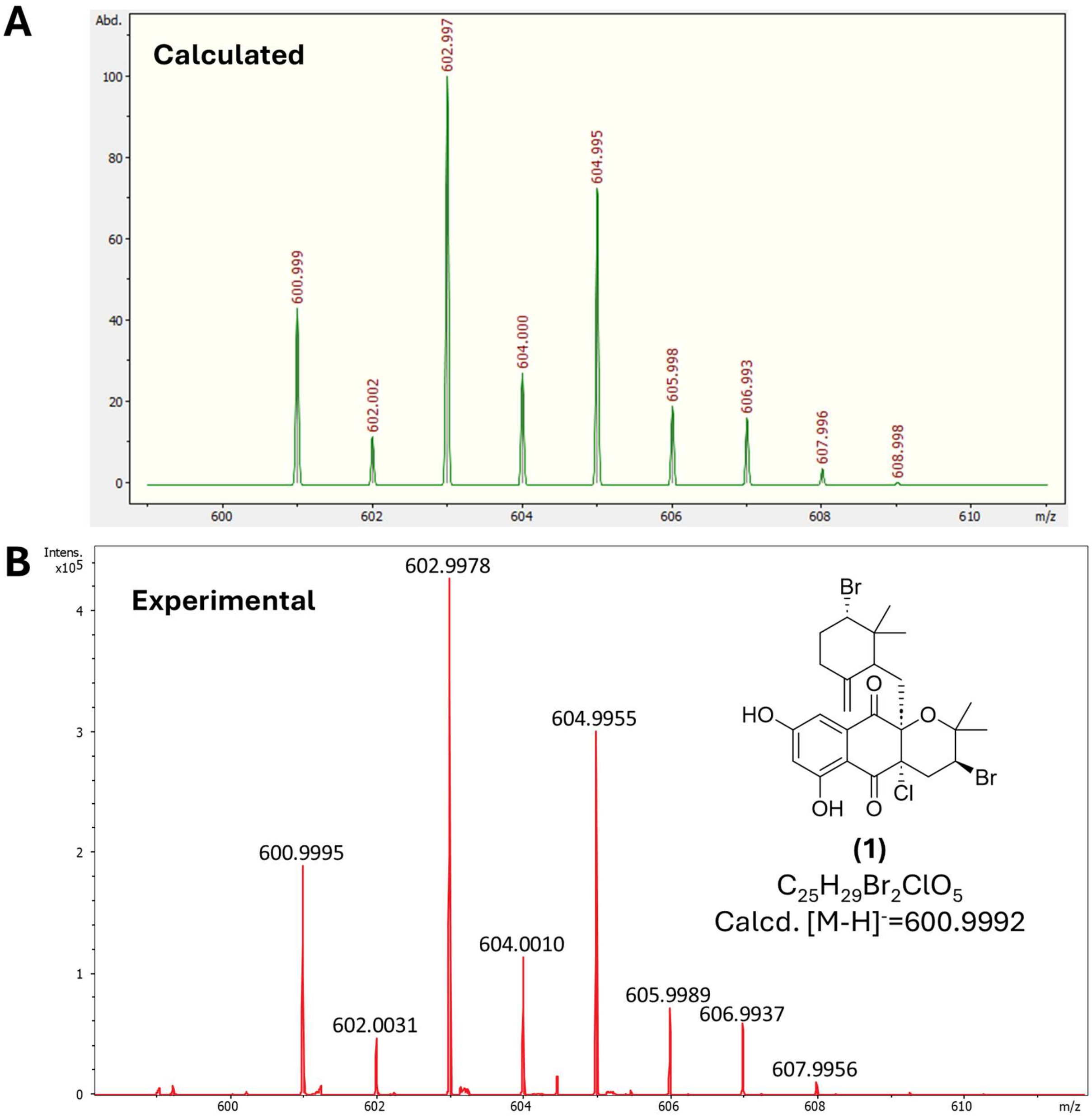


**Figure S3** MALDI-MS[1] spectrum and isotopic distribution of napyradiomycin B8 (**1**). **A)** Isotopic distribution calculated for $[M\text{-}H]^-$ of $C_{25}H_{29}Br_2ClO_5$ using Bruker's IsotopePattern tool. The formula and $[M\text{-}H]^-$ were calculated using $^{79}Br$ and $^{35}Cl$. **B)** Experimental MS[1] spectrum of compound **1**. The isotopic distribution of the experimental spectrum is highly consistent with that predicted for the $[M\text{-}H]^-$ of $C_{25}H_{29}Br_2ClO_5$.

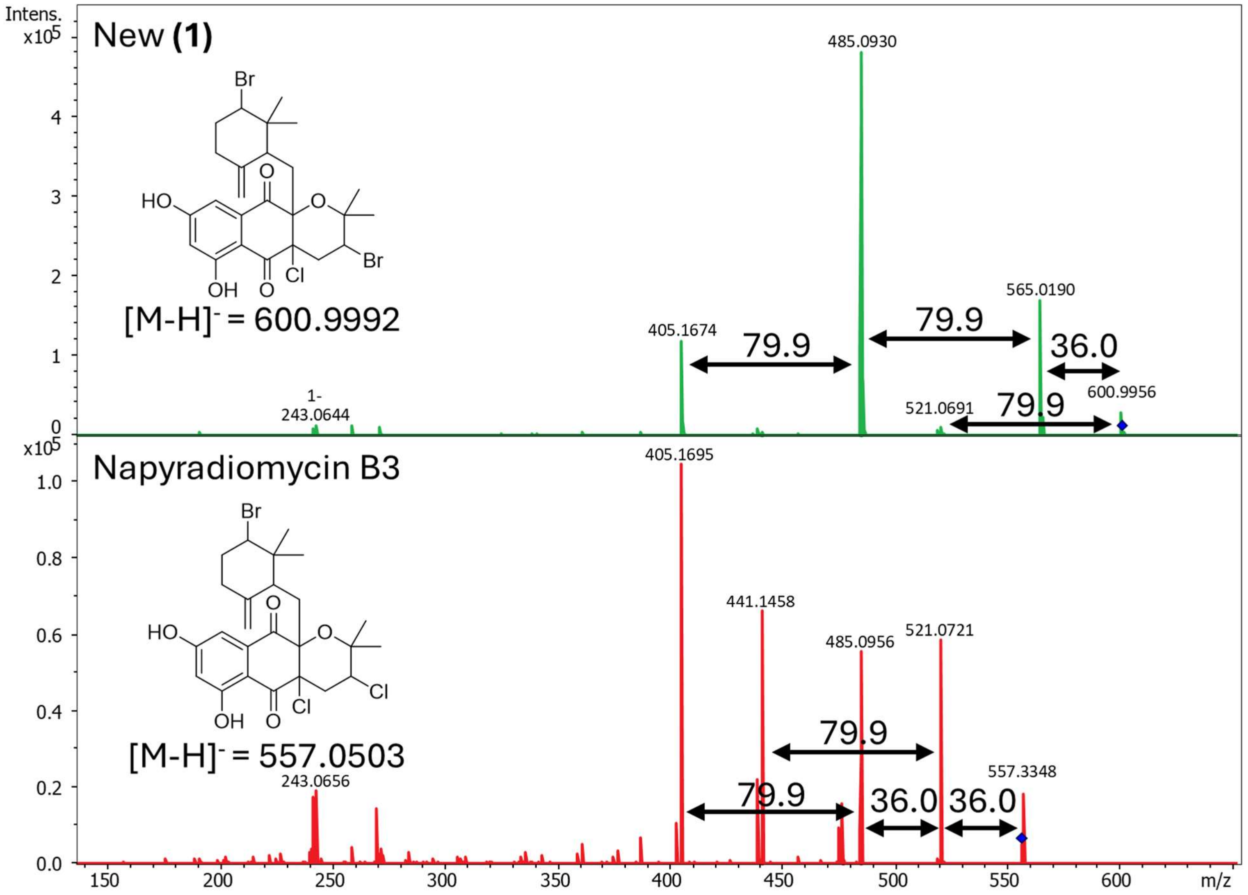


**Figure S4** MALDI-MS/MS spectra of compound **1** (top), versus that of napyradiomycin B3 standard (bottom). Loss of 79.9 Da corresponds to consecutive loss of HBr, while loss of 36.0 Da corresponds to consecutive loss of HCl from the core napyradiomycin B scaffold.

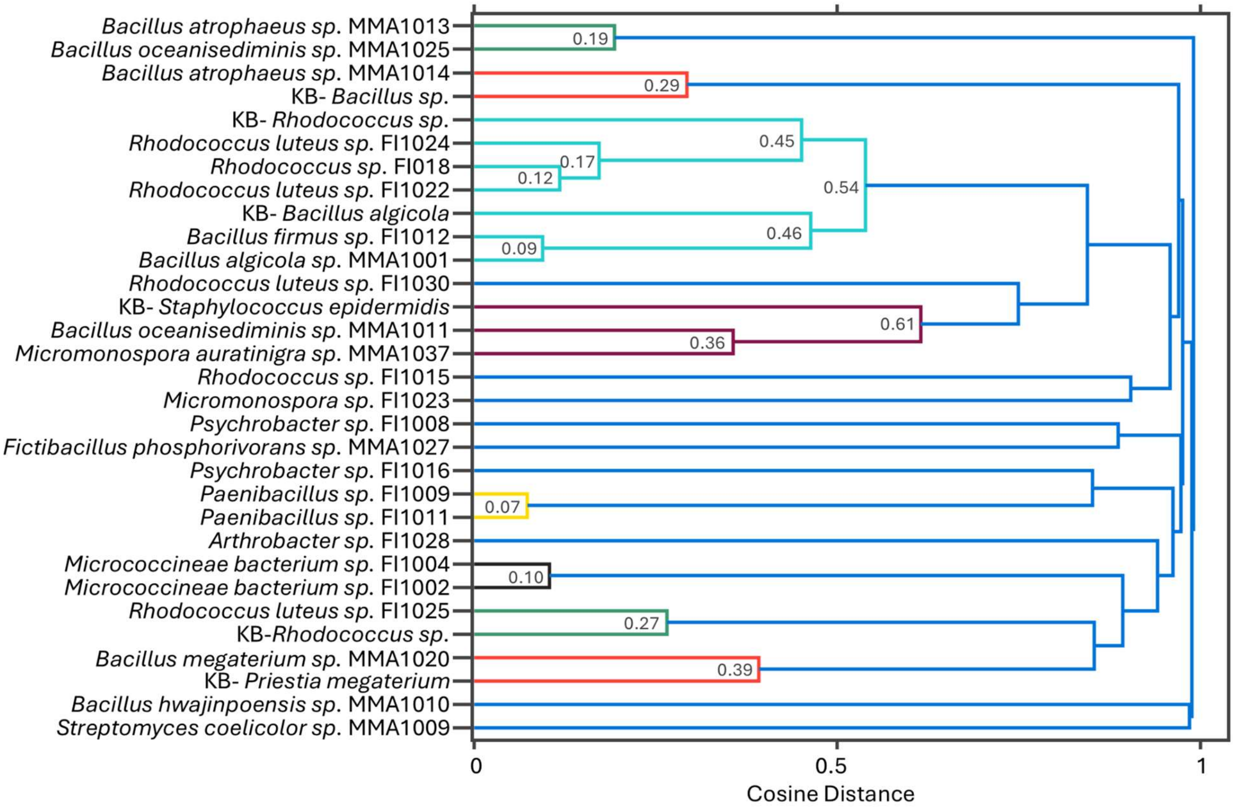


**Figure S5** Protein spectral dendrogram of 25 commensal microbial strains previously isolated from marine vertebrate intestines.[1,2] All strains were previously identified using 16s rRNA sequencing. Taxonomic clustering between the analyzed strains and within the IDBac knowledge base appears to be highly consistent with 16s rRNA taxonomy.

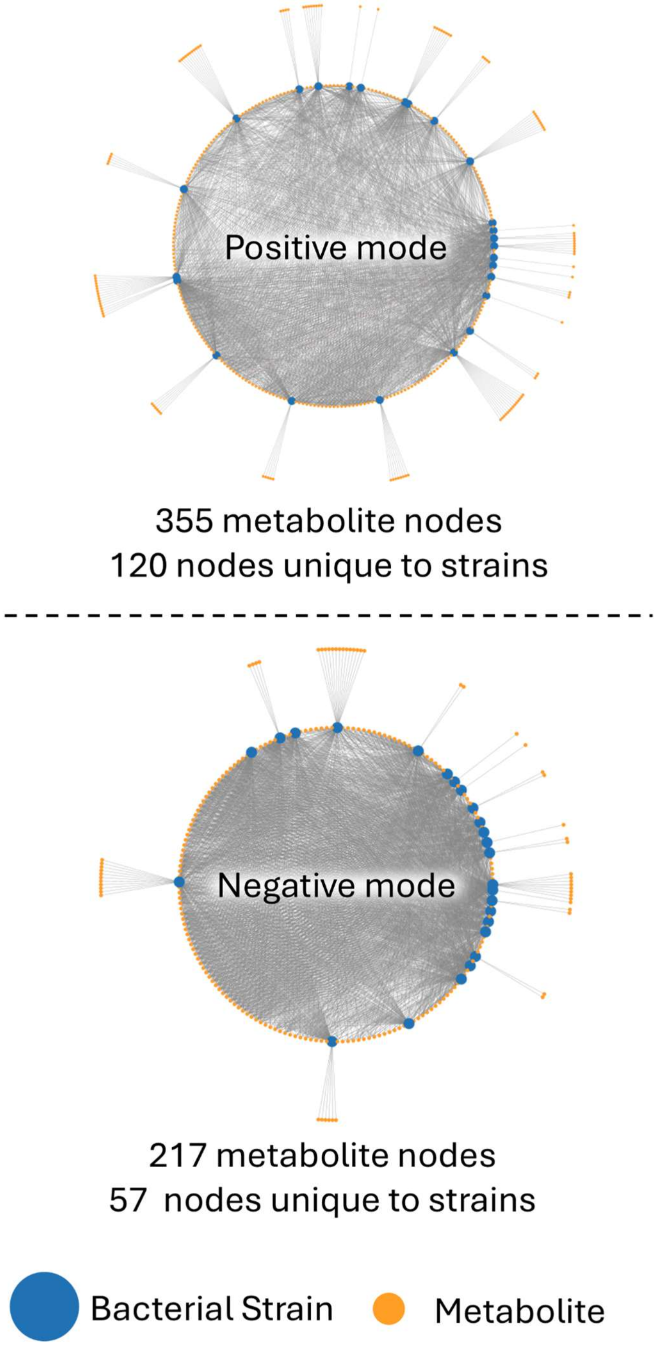


**Figure S6** Positive and negative mode metabolite association networks of 25 commensal microbial strains previously isolated from marine vertebrate intestines.[1,2]

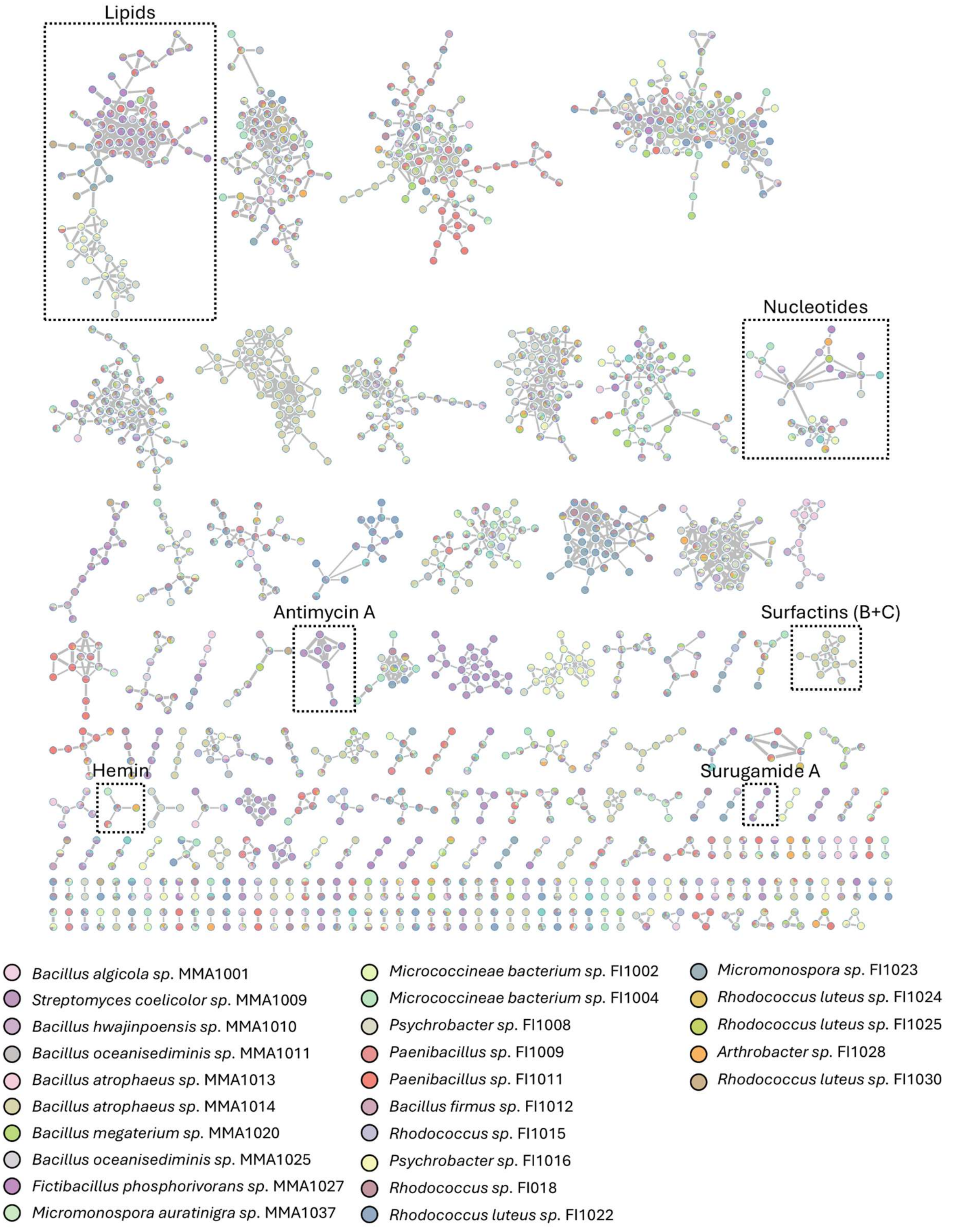


**Figure S7** Combined positive and negative mode molecular network composed of MALDI-MS/MS data collected directly from colonies of 25 commensal bacteria. Nodes are colored based on individual bacterial strains. Relevant node counts for individual strains are listed in **Table 1** in the main text.

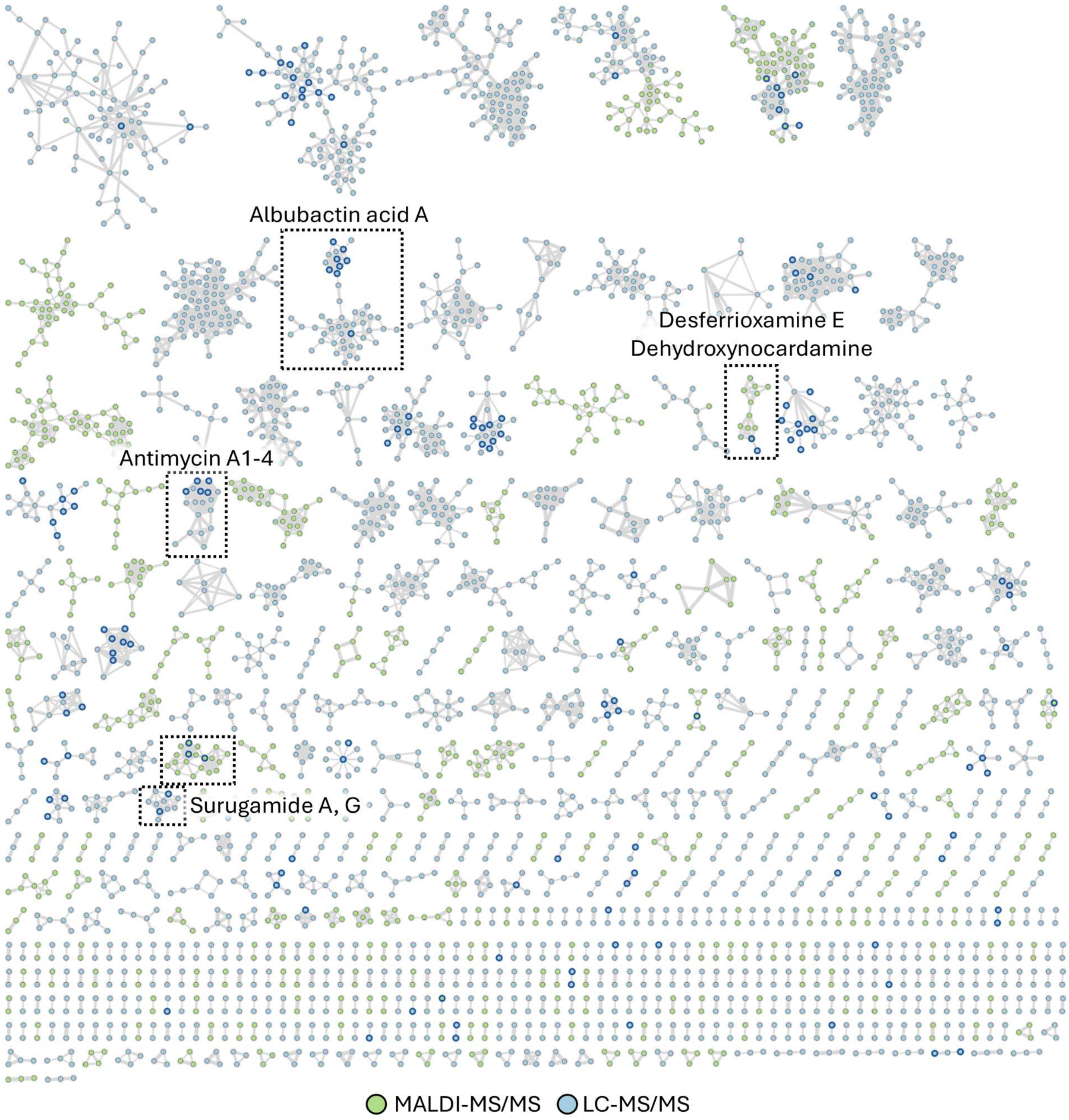


**Figure S8** Combined MALDI- and LC-MS/MS molecular network of SPE fractions A-F of *Streptomyces coelicolor sp.* MMA1009. Dotted boxes are drawn around the molecular families highlighted in **Figure 5** of the main text. Nodes with dark blue borders indicate GNPS2 library hits.

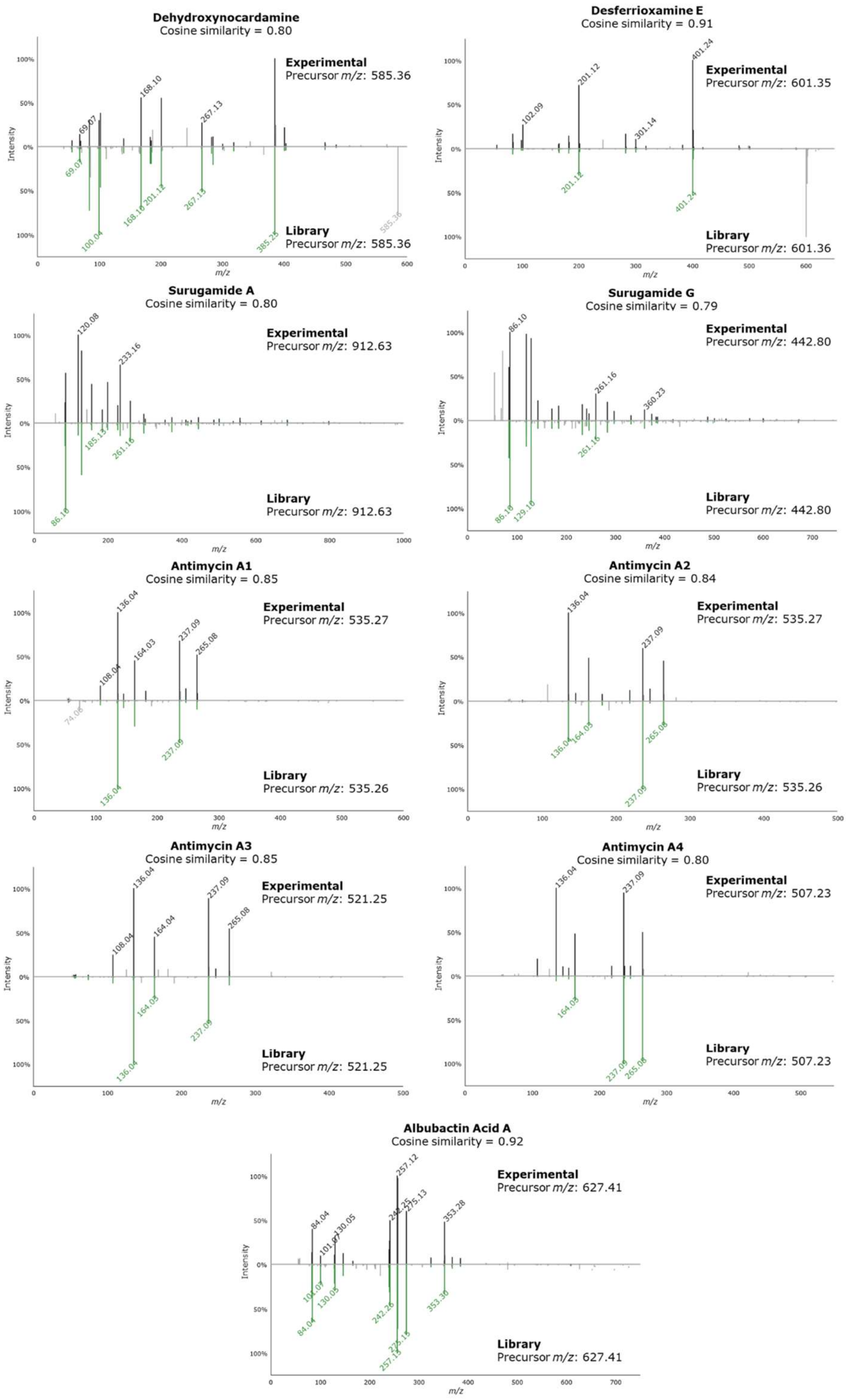


**Figure S9** Mirror plots of combined MALDI- and LC-MS/MS spectra from molecular network of SPE fractions A-F of *Streptomyces coelicolor sp.* MMA1009 as highlighted in **Figure 5.**

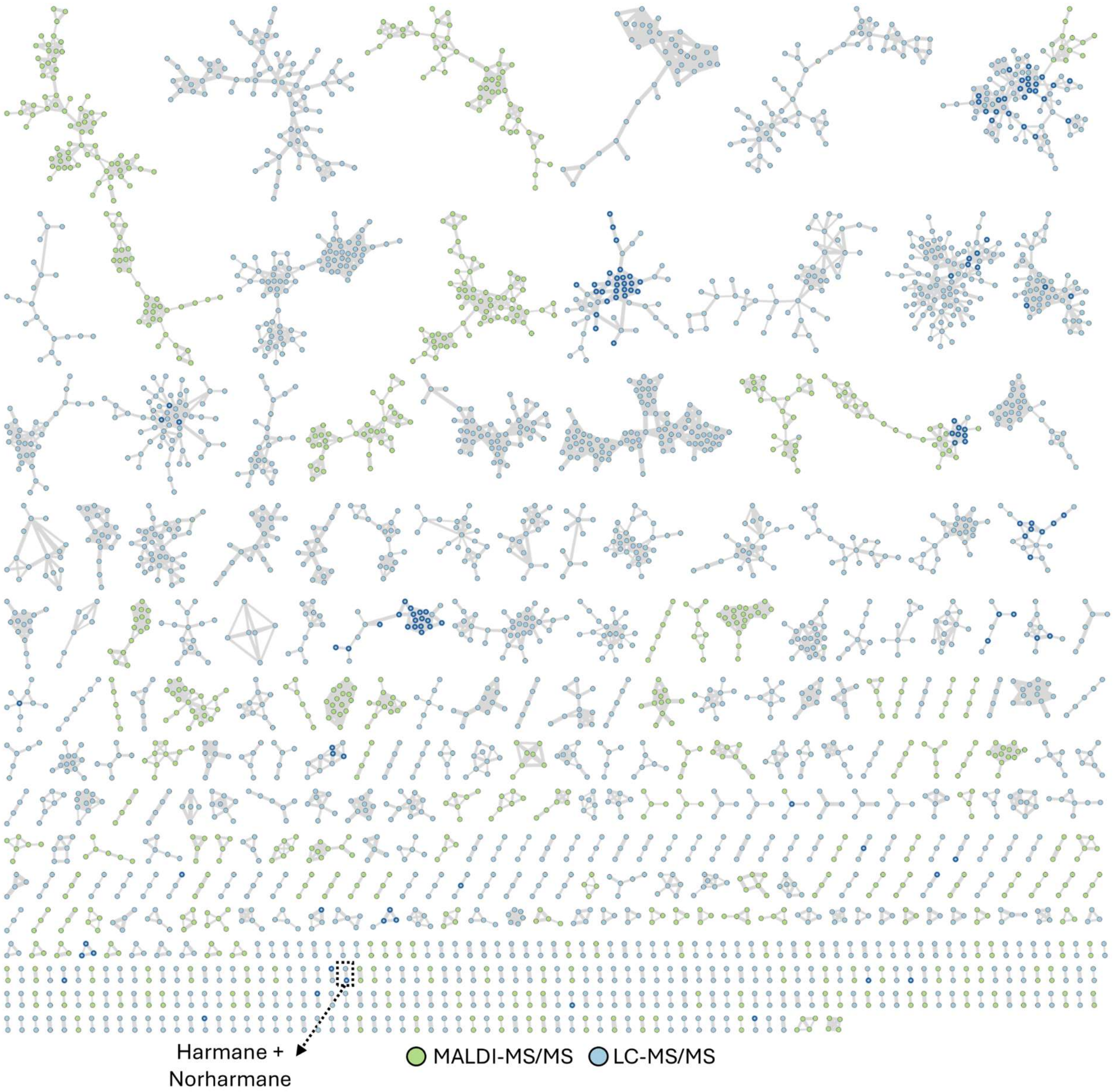


**Figure S10** Combined MALDI- and LC-MS/MS molecular network of SPE fractions A-F of *Psychrobacter sp.* FI1016. Nodes with dark blue borders indicate GNPS2 library hits.

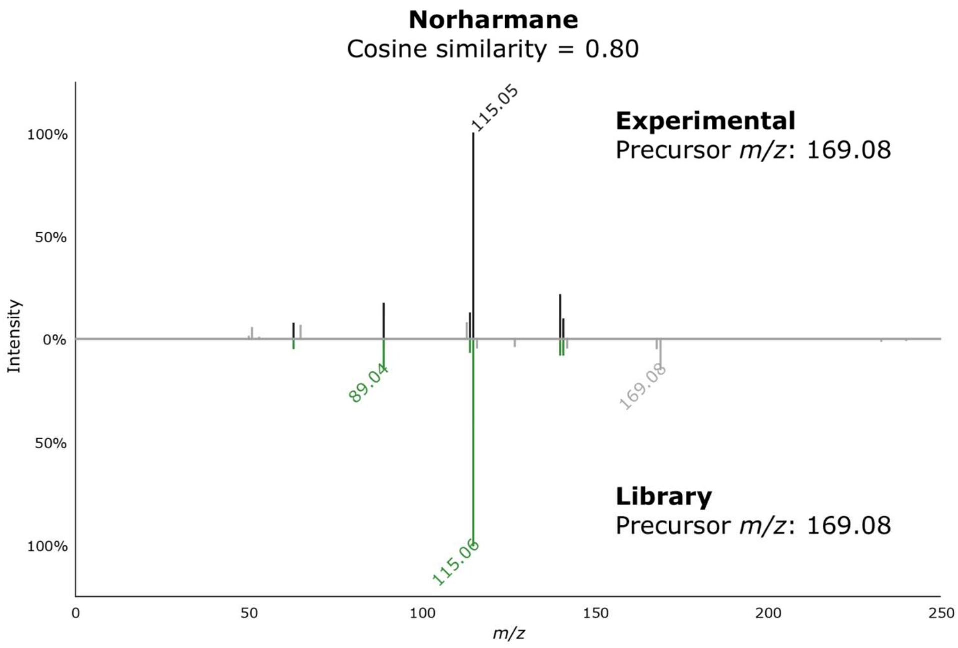


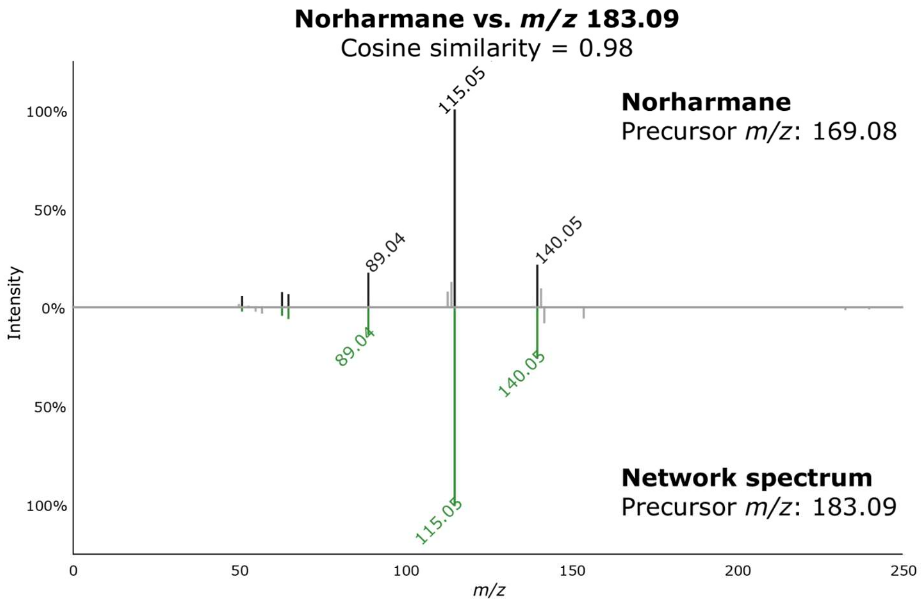


**Figure S11** MS/MS spectrum of *m/z* 169.08 against the GNPS2 library spectrum of norharmane (top) and MS/MS spectrum of *m/z* 169.08 vs *m/z* 183.09 (bottom), which was later determined to be harmane via NMR analysis.

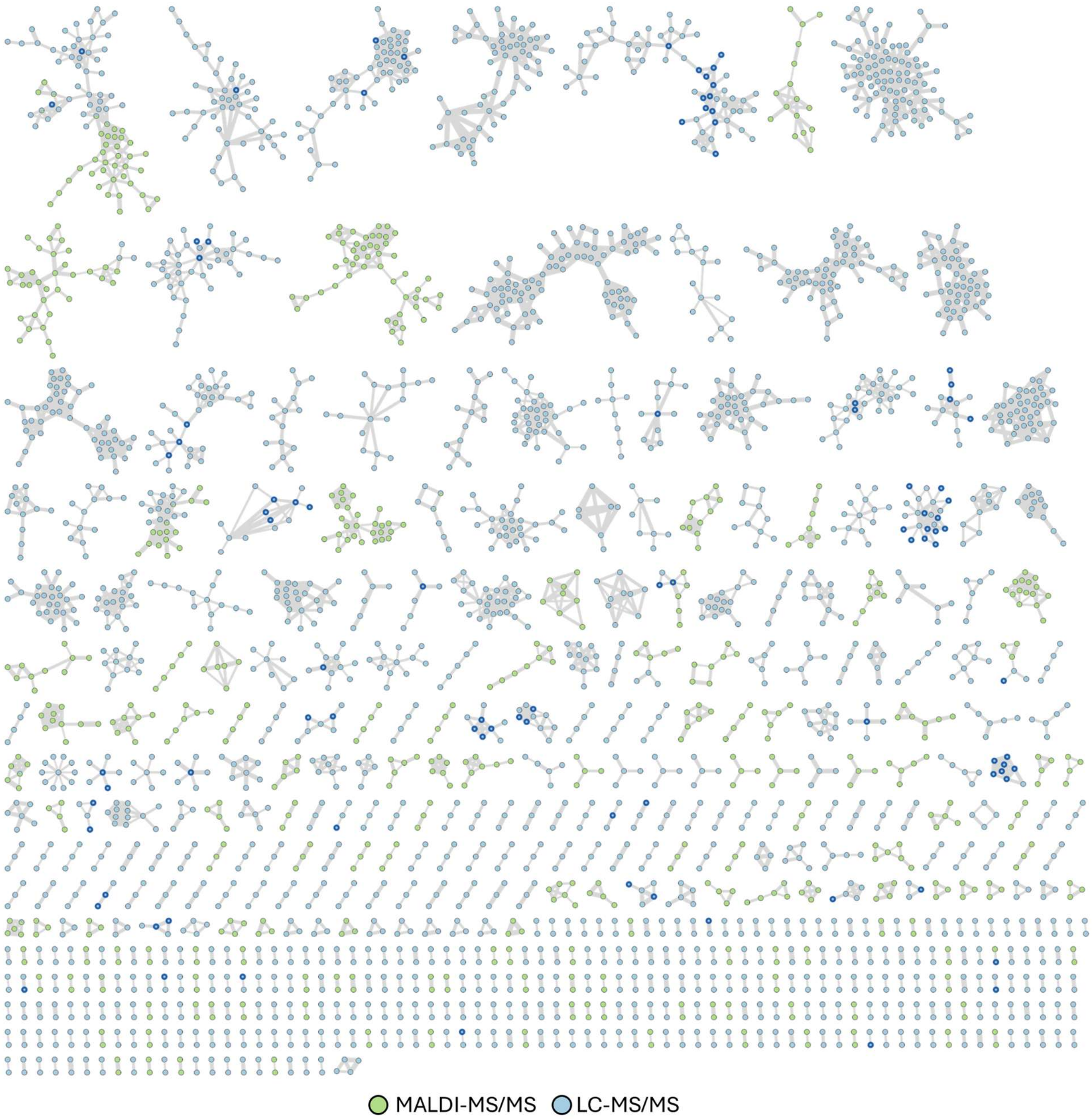


**Figure S12** Combined MALDI- and LC-MS/MS molecular network of SPE fractions A-F of *Micromonospora sp.* FI1023. Nodes with dark blue borders indicate GNPS2 library hits.

**NMR Data**

Labeling scheme used for napyradiomycin B compounds based on the original isolation paper.[3]

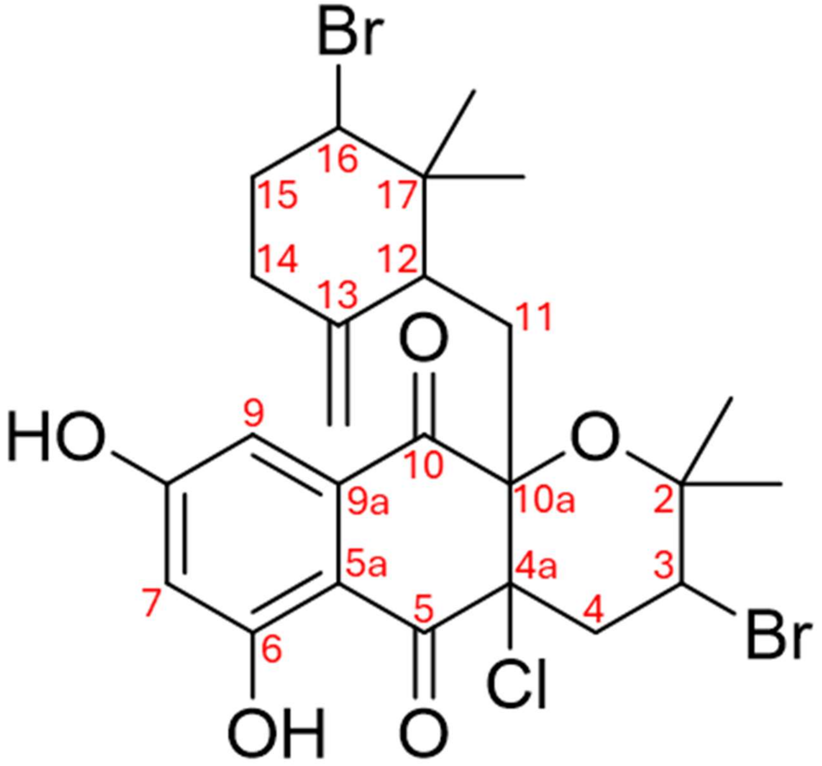

**Table S5** $^1H$ NMR comparison of the isolated napyradiomycin B8 (**1**) with the most recently reported $^1H$ NMR data of napyradiomycin B1.[4]

| Assignment | Napyradiomycin B1 | Napyradiomycin B8 (1) |
|---|---|---|
| 2-$CH_3$ | 1.38, s | 1.39, s |
| 2-$CH_3$ | 1.19, s | 1.27, s |
| 3 | 4.45, dd<br>(12.0, 4.0) | 4.57, dd<br>(12.2, 3.9) |
| 4 | 2.52, dd<br>(14.1, 3.9)<br>2.35, dd<br>(14.1, 12.0) | 2.61, dd<br>(14.0, 3.9)<br>2.49, dd<br>(13.9, 12.4) |
| 6-OH | 12.04, s | 12.03, s |
| 7 | 6.73, d<br>(2.5) | 6.71, br s |
| 8-OH | 7.10, br s | Not observed |
| 9 | 7.14, d<br>(2.5) | 7.11, d<br>(2.3) |
| 11 | 2.64, dd<br>(15.5, 8.7)<br>1.62, d<br>(15.2) | 2.66, dd<br>(15.7, 8.7)<br>1.62, (COSY) |
| 12 | 1.99, d<br>(8.6) | 2.04, d<br>(8.8) |
| 13-$CH_2$ | 4.78, d<br>(1.4) | 4.77, d<br>(7.2) |
| 14 | 2.24, ddd<br>(13.2, 4.2, 4.2)<br>1.94, ddd<br>(13.3, 13.3, 4.9) | 2.19*, m<br><br>1.93, m |
| 15 | 2.03, dddd<br>(12.7, 4.6, 3.5, 3.5)<br>1.75-1.68, m | 2.19*, m<br><br>Buried |
| 16 | 3.80, dd<br>(11.6, 4.4) | 4.05, dd<br>(11.1, 4.4) |
| 17-$CH_3$ | 0.58, s | 0.63, s |
| 17-$CH_3$ | 0.71, s | 0.73, s |

*Overlapping signals

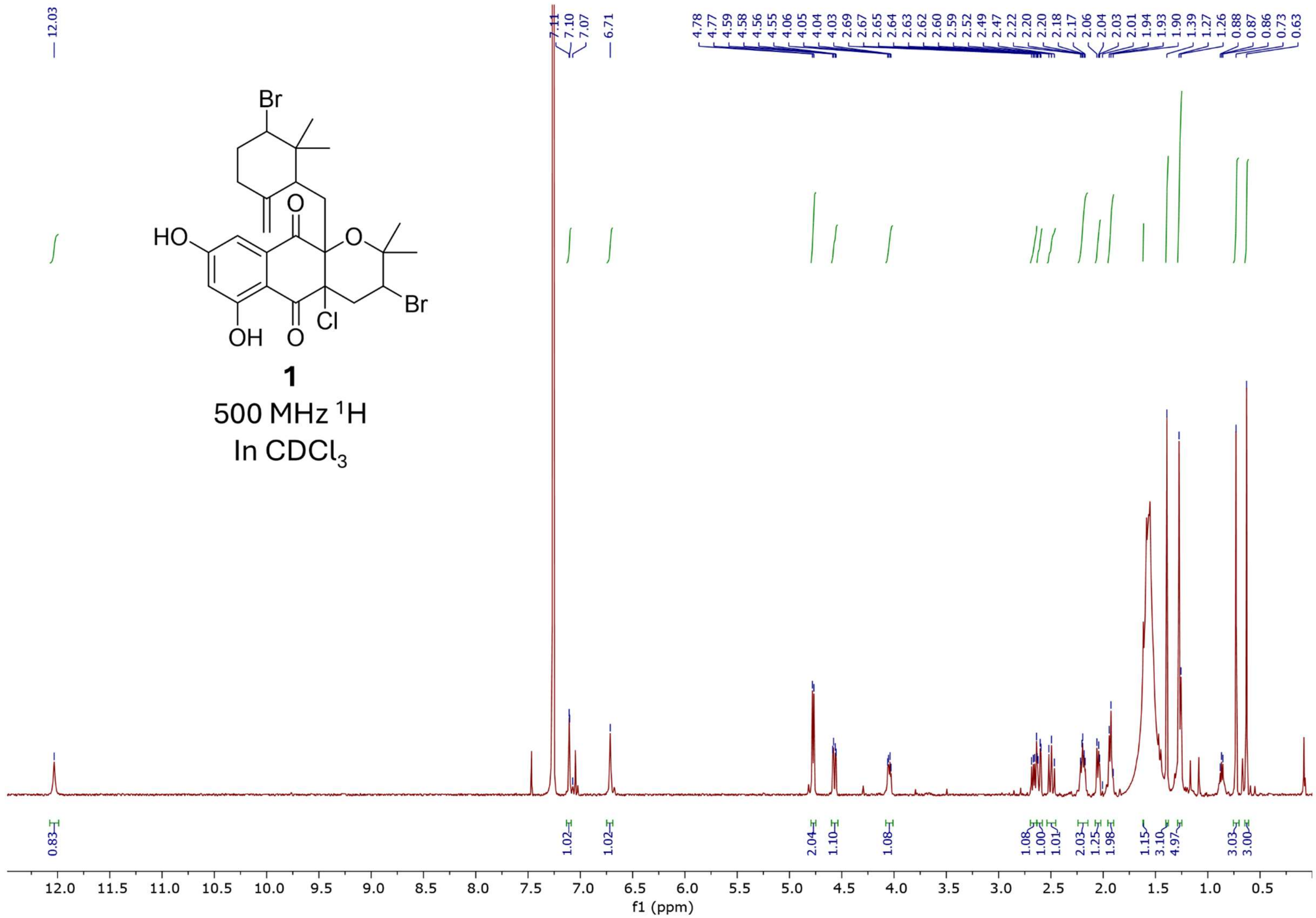


**Figure S13** $^{1}$H NMR spectrum of compound **1** in $CDCl_3$, 500 MHz.

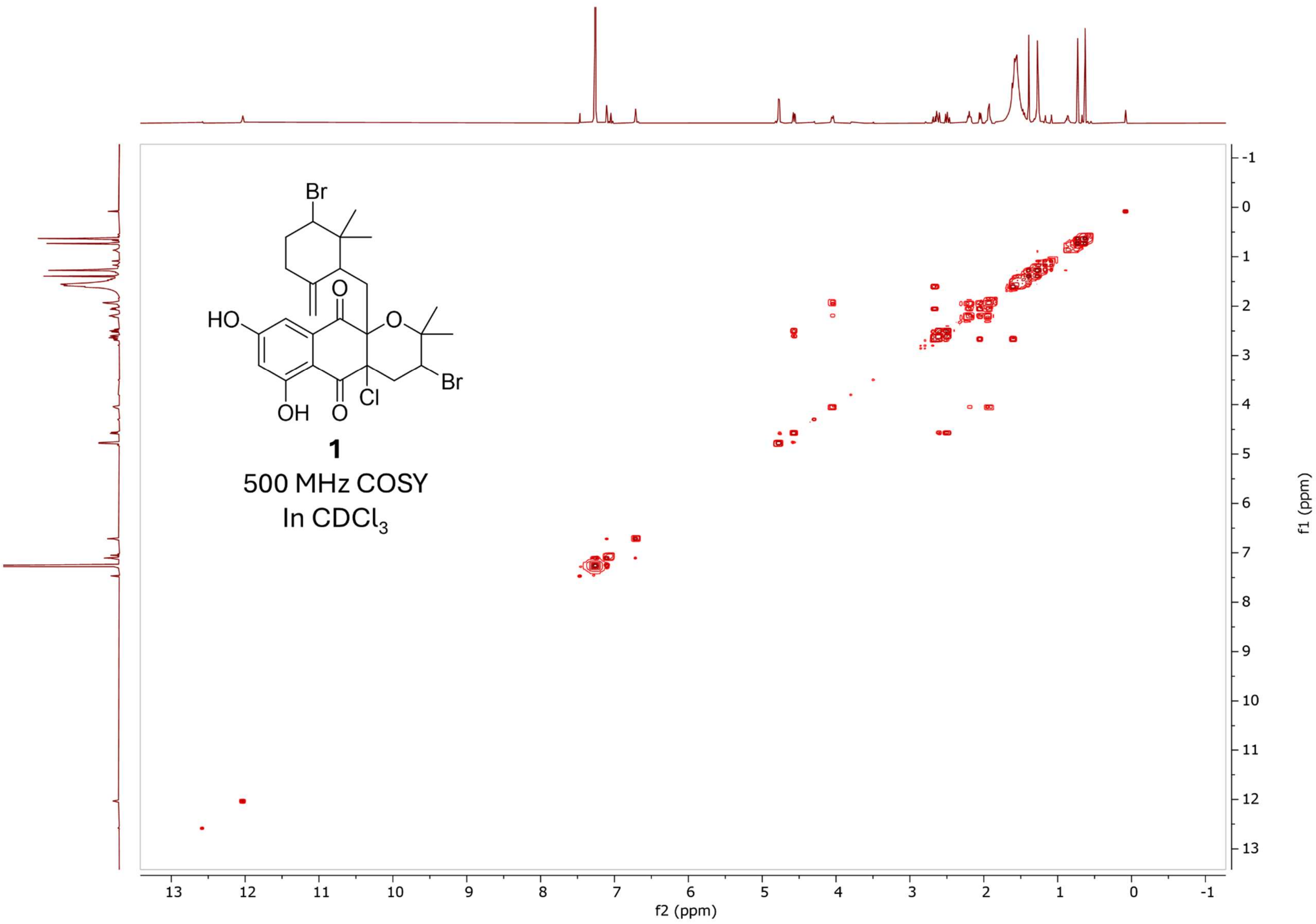


**Figure S14** $^1$H-$^1$H Gradient COSY spectrum of compound **1** in $CDCl_3$, 500MHz.

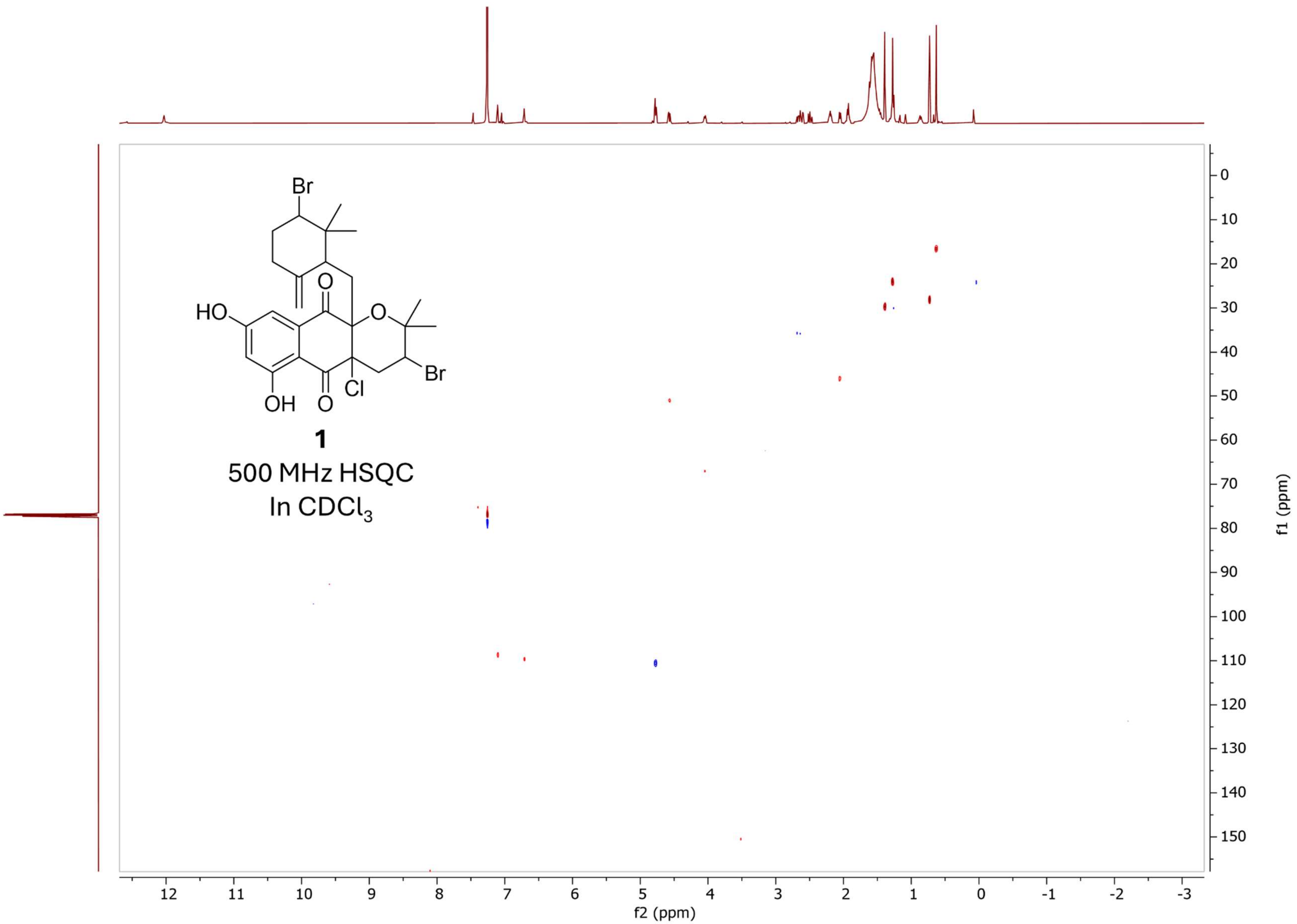


**Figure S15** Multiplicity-edited HSQC of compound **1** in $CDCl_3$, 500 MHz.

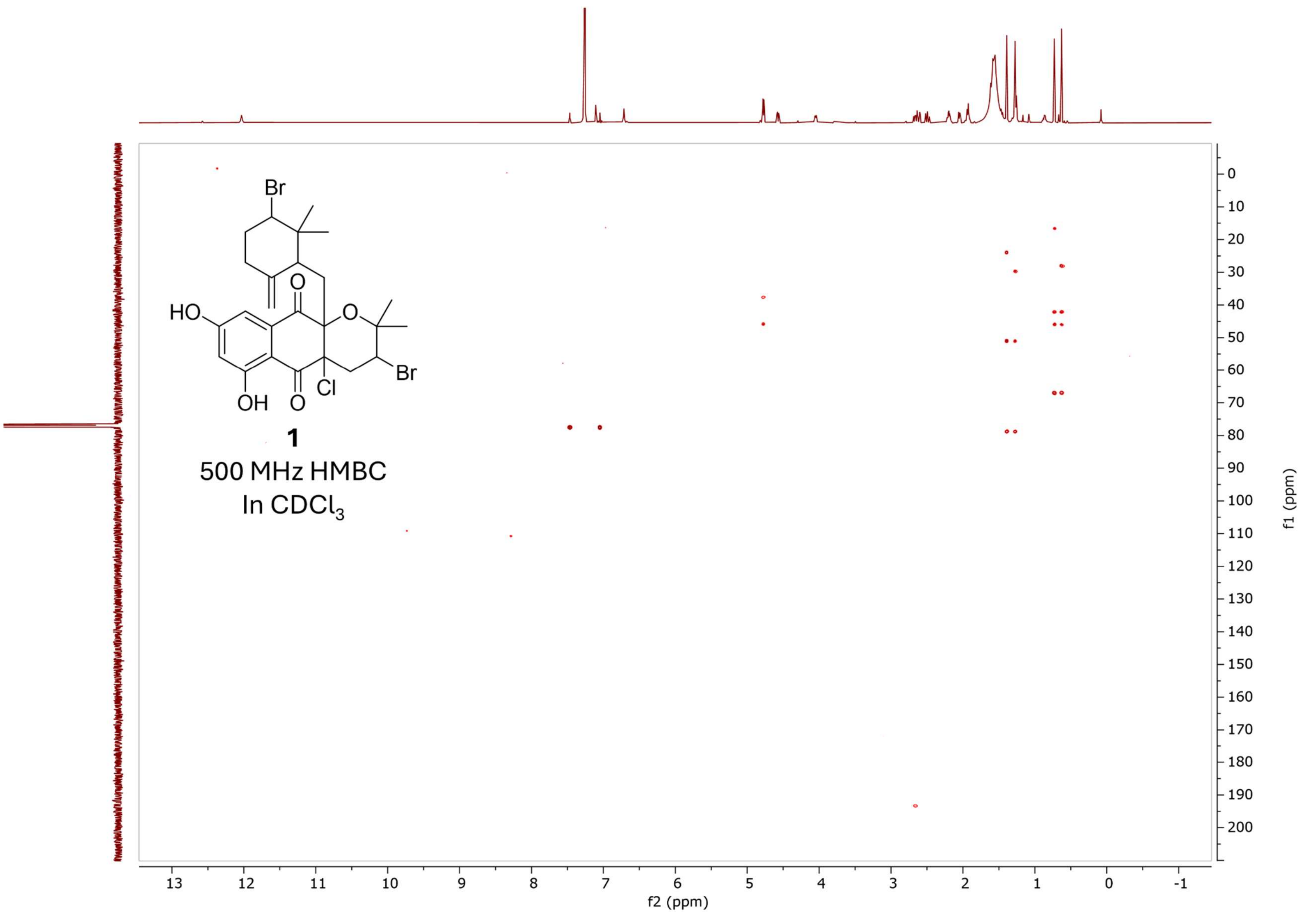


**Figure S16** HMBC spectrum of compound **1** in $CDCl_3$, 500 MHz.

Labeling scheme used for harmane and norharmane based on that outlined by Seki et al.[5]

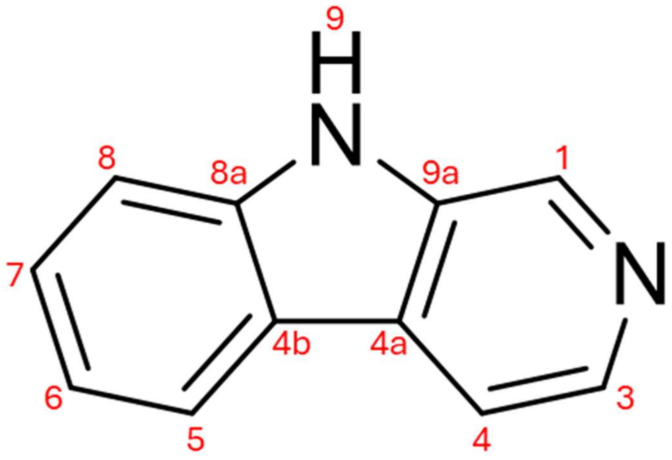


Norharmane (2)

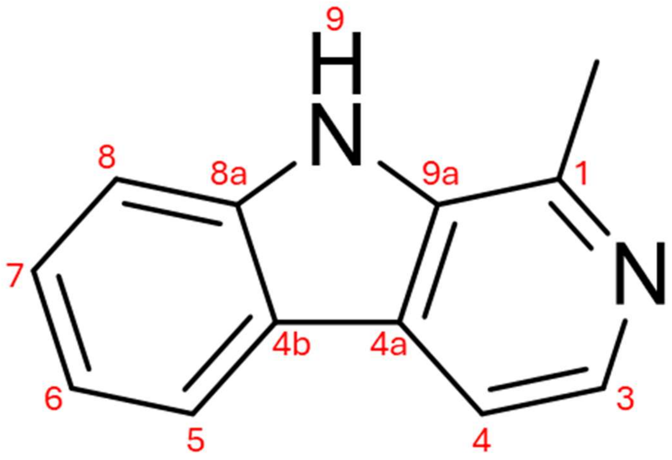


Harmane (3)

**Table S6** $^1H$ NMR comparison of the isolated norharmane (**2**) with the values reported by Siderhurst et al. in $CD_3OD$.[6]

| Assignment | Norharmane (Siderhurst et al.[6]) 300 MHz | Norharmane (**2**) 800 MHz |
|---|---|---|
| 1 | 8.79 s | 8.82 s |
| 3 | 8.29 d (5.4) | 8.31 (5.3) |
| 4 | 8.09 d (5.4) | 8.12 (5.3) |
| 5* | 7.58 m | 7.59 m |
| 6 | 7.28 m | 7.29 ddd (7.9, 5.9 , 2.0 ) |
| 7* | 7.58 m | 7.59 m |
| 8 | 8.2 d (7.9) | 8.2 d (7.9) |
| 9-*N*-H | Not observed | Not observed |

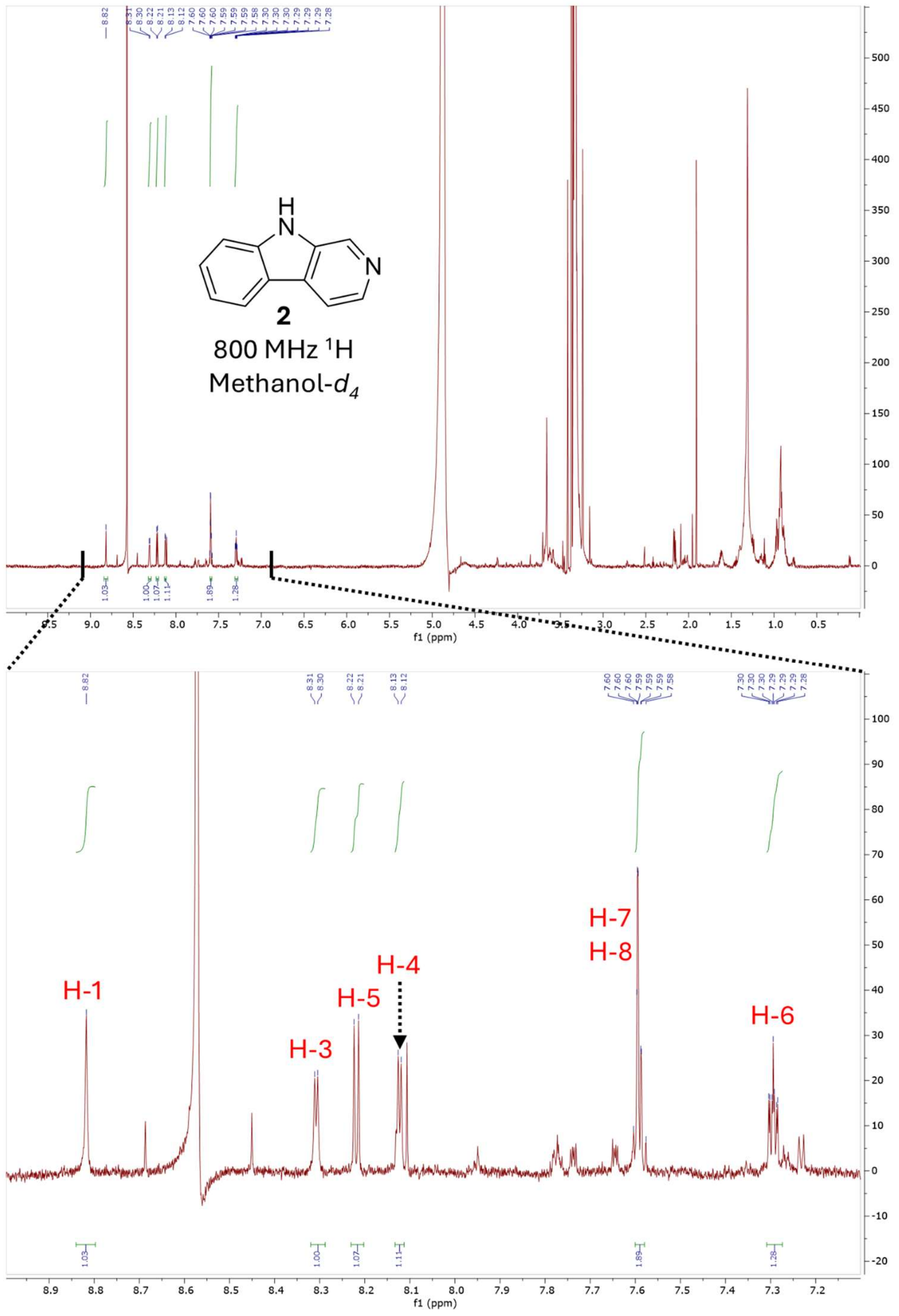


**Figure S17** $^{1}$H NMR spectrum of norharmane (**2**) in $CD_3OD$, 800 MHz.

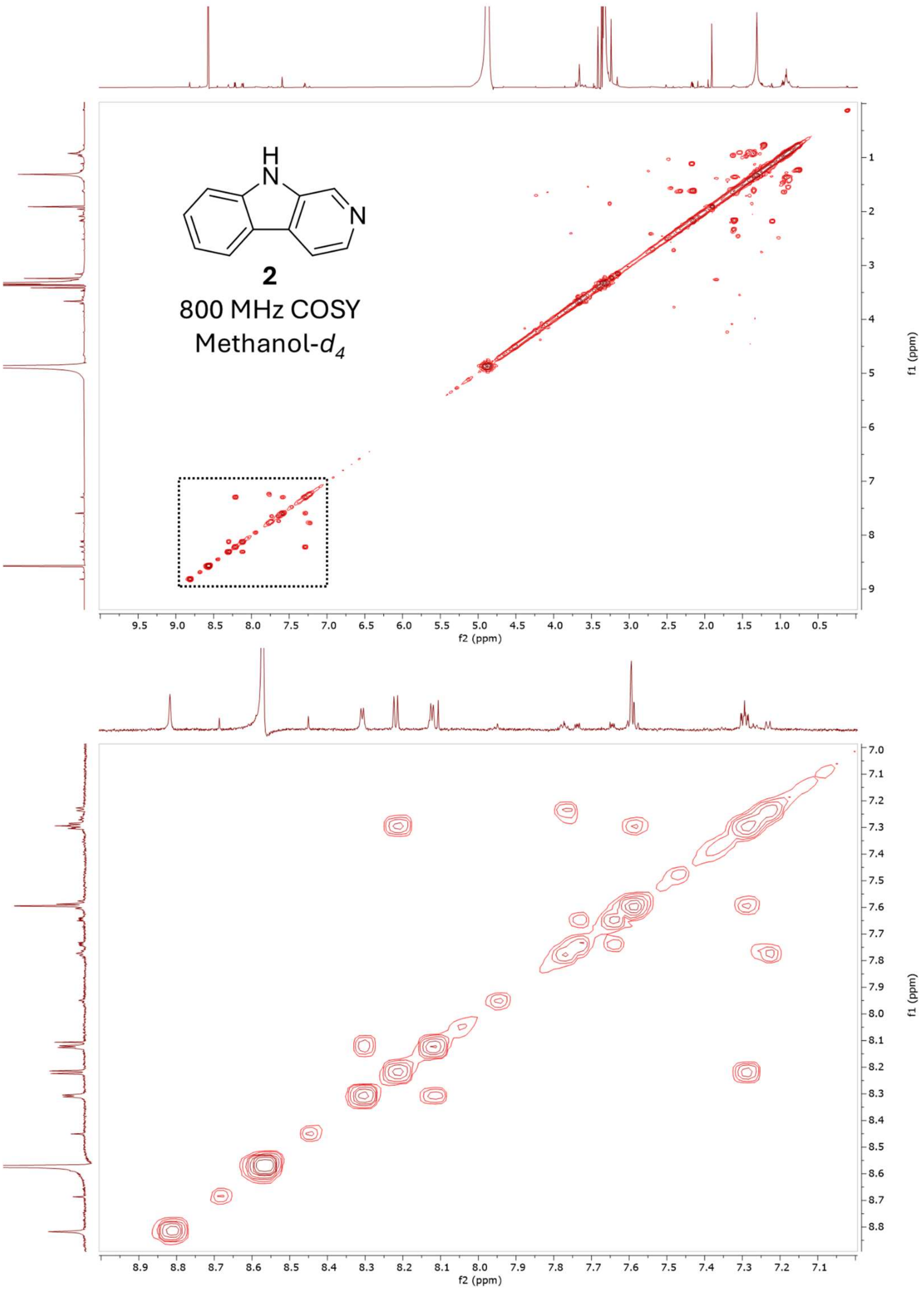


**Figure S18** COSY spectrum of norharmane (**2**) in $CD_3OD$, 800 MHz.

**Table S7** $^1H$ NMR comparison of the isolated harmane (**3**) with the values reported by Seki et al. in $CD_3OD$.[5]

| Assignment | Harmane (Seki et al.[5]) | Harmane (**3**) 800 MHz |
|---|---|---|
| 1-$CH_3$ | 2.80 s | 2.83 s |
| 3 | 8.15 d (5.5) | 8.18 d (4.8) |
| 4 | 7.90 d (4.5) | 7.95 d (5.5) |
| 5 | 8.13 dd (8.2, 0.74) | 8.17 d (4.3, 0.7) |
| 6 | 7.24 td (7.4, 1.1) | 7.27 td (6.2, 0.9) |
| 7 | 7.53 td (6.9, 1.1) | 7.56 td (6.9,1.2) |
| 8 | 7.58 dd (8.2, 0.7) | 7.61 d (8.2, 0.7) |
| 9-*N*-H | Not observed | Not observed |

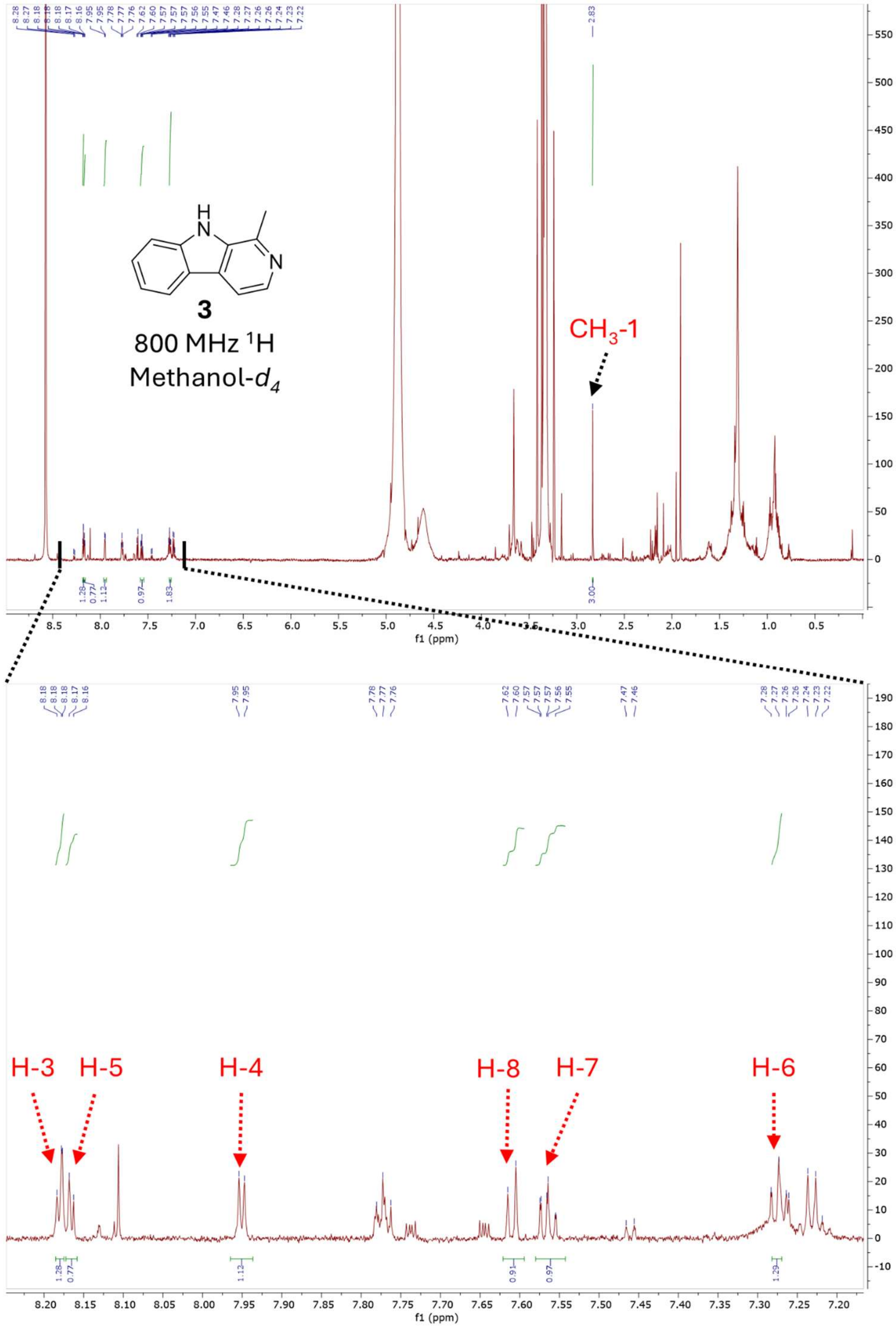


**Figure S19** $^{1}$H NMR spectrum of harmane (**3**) in $CD_3OD$, 800 MHz.

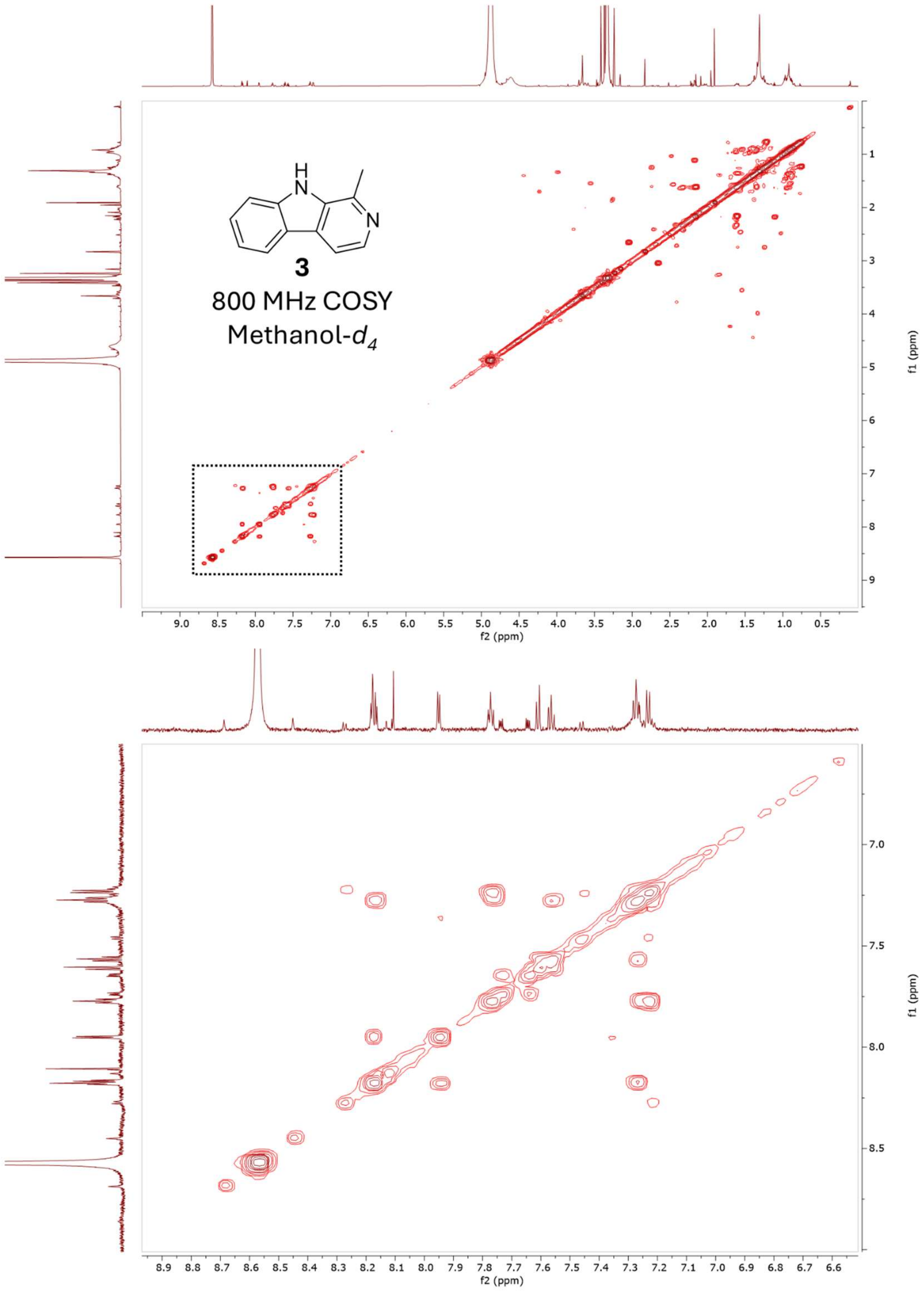


**Figure S20** COSY spectrum of harmane (**3**) in $CD_3OD$, 800 MHz.

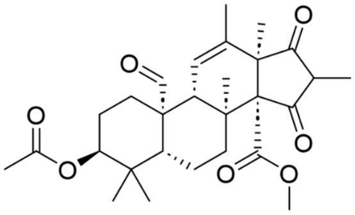

**Andrastin A**
$C_{28}H_{38}O_7$
$[M+H]^{+}$= 487.2690

**Andrastin A 15 eV**

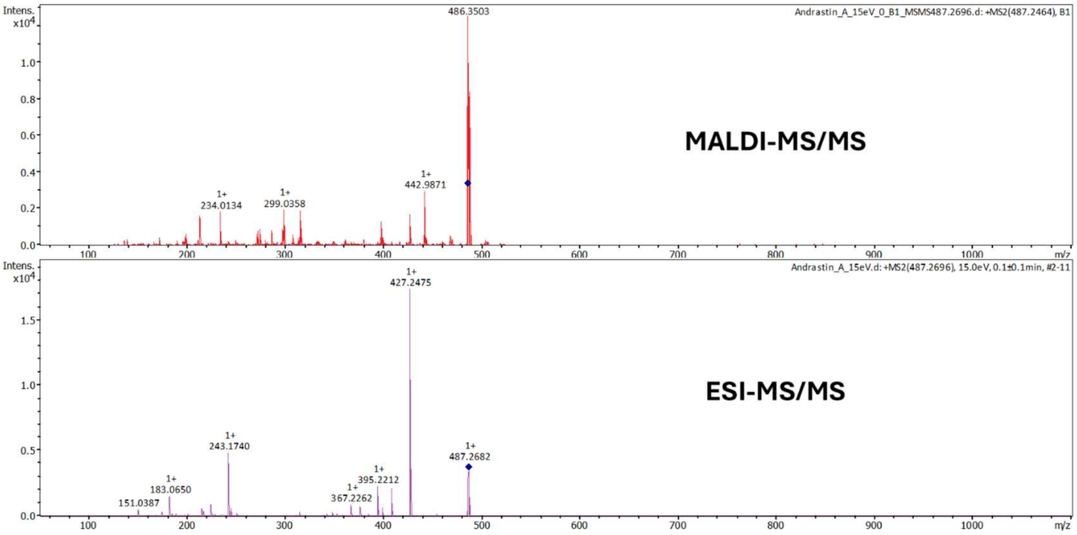


**Andrastin A 25 eV**

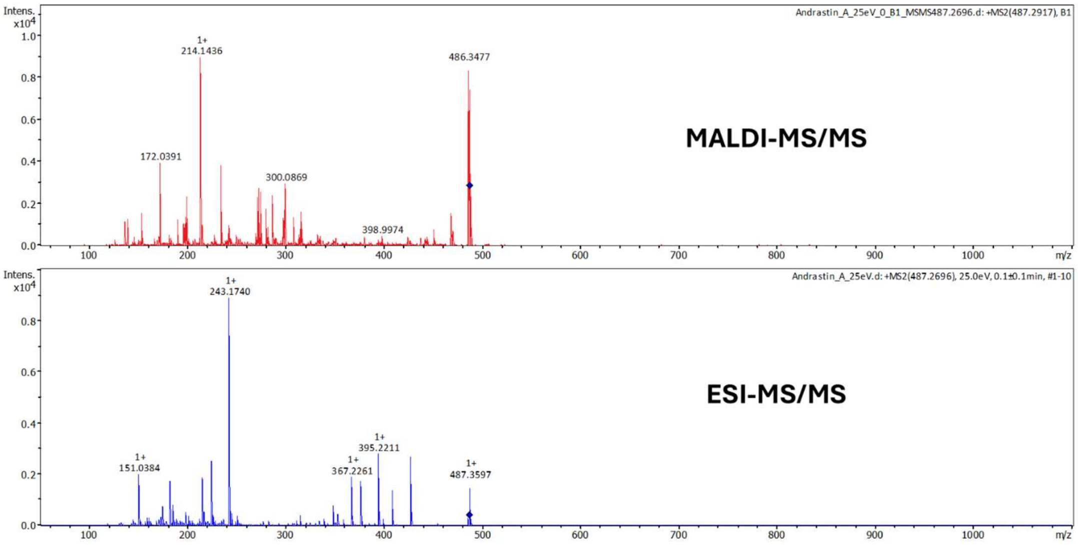


**Figure S21** Comparison of fragmentation of andrastin A between MALDI- and ESI-MS/MS at 15 and 25 eV. The fragmentation spectra are different between the two ionization modalities, despite being activated via collision induced dissociation (CID) on the same instrument (the timsTOF fleX).

**Supporting Information References:**
